# Pictorial balance, a bottom-up neuro-aesthetic property mediating attention and eye movements, explains the feeling of unity and harmony in pictures. A primitive visual operating system determines balance and shows how the world was first visually organized using luminance and movement

**DOI:** 10.1101/2020.05.26.104687

**Authors:** David Corwin

## Abstract

A computer model of a perfectly balanced picture is created. Analysis allowed the creation of an algorithm for calculating balance based only on the quadrant luminance. A study shows a correlation between observers’ ability to determine relative balance. The algorithm is a potential operating system for primitive organisms to identify and assess threat potential of other organisms. From this perspective an ecological description of the study explains the correlation in terms of causation. The luminance information used to determine balance originates in peripheral vision and follows the tectopulvinar pathway. It is in competition for attention with foveal information and suppresses magnocellular information that both use the geniculostriate pathway. In an unbalance picture this inhibits peripheral retinal information and saccadic movement. This is the basis of the 20^th^ century force-field theory of pictorial balance. Seeing the picture as a whole using peripheral vision evoking feelings of unity and being able to move smoothly through it evoking feelings of harmony is the basis of the neuro-aesthetic effects of a picture.

Pictorial balance is used to explain why some paintings evoke the aesthetic feelings of unity and harmony. The original unsubstantiated concept of pictorial balance as a center of mass effect, i.e. the feeling that somehow a person on the left seems to be balanced by a tree on the right, originated at the beginning of the 20^th^ century. This study of balance starts with an elusive and unnamed pictorial effect known to painters since the 17^th^ century that evokes feelings of harmony and unity. In such pictures the image seems to be perceived as a whole without the desire to fixate on individually depicted objects. The author thought that this effect indicated that the picture was in a state of perfect balance. Computer modeling found that such a picture should have bilateral luminance symmetry with a slightly lighter lower half, and that with respect to balance the eye could not distinguish the picture from its white frame. It was proposed that a pictures balance could be calculated as a property of a moving luminous object by an algorithm to identify and follow other organisms. A study was done in which observers viewed pairs of identical pictures in different frames and were asked to say if they appeared different. It was found that the extent to which the picture pairs were seen as different is inversely correlated with balance as calculated from the algorithm. The results are consistent with the pictures being seen on a low level as living organisms. Balance is perceived with peripheral vision; a picture is seen as an object giving it a feeling unity. As a picture becomes unbalanced, the eye will look in the picture at what is depicted. The conflict between these two ways creates tension that explains why earlier researchers postulated the existence of force fields within the pictorial plane.

## Introduction

Pictures are flat, bounded images that correspond to nothing found in nature or mental imagery, including the borderless images in prehistoric caves ^1^. They are a very recent invention requiring the technology to create flat surfaces dating from perhaps the fourth millennium BC, and differ from images without borders. Borders evoke the percept of balance: the parts relate to both the center and the border ^2 p.10^. Balance is both a bottom-up percept and a top-down aesthetic concept taught in schools. This paper is concerned with the bottom-up aesthetic perception – the pre-emotional, pre-cognitive pathway from the retina to the primary visual cortex as Molnar postulated ^3^.

It has been stated that “for centuries artists and writers on Western art have asserted that balance is the primary design principle for unifying structural elements of a composition into a cohesive narrative statement.” A picture has been considered balanced “if all its elements are arranged in such a way that their perceptual forces are in equilibrium about a fulcrum and if the fulcrum coincides with the center of the picture.”^4^ Objects, size, color, intrinsic interest, positions, and apparent depth have been proposed as contributing to this balance and the aesthetic qualities of unity and harmony if they are balanced around the geometric center as calculated by a center-of-mass type of equation as if a picture evokes a force field.^2,4–6^ Rather than being a timeless observation, it is an early 20^th^-century concept promoted initially by Henry Rankin Poore, a painter. However, he was most likely trying to explain the effect of low-level balance that I discuss in this paper in that he wrote:

> “The whole of the pictorial interest may be on one side of a picture and the other side be practically useless as far as picturesqueness or story-telling opportunity is concerned, but which finds its reason for existing in the balance, and that alone.”

Nothing in this “useless” side has the pictorial qualities described above, but for him they create in a particular picture a feeling that he ascribes to the feeling of being balanced.^5^ Several studies have unsuccessfully attempted to confirm this method of calculating balance.^7–9^ However, both Hubert Locher and Thomas Puttfarken give comprehensive historically discussions of European theories of pictorial aesthetics and composition between the 14^th^ and the late 19^th^ century in which balance is not mentioned.^10,11^ This judgment of balance is inherently a modern top-down idea taught in schools. A Google Ngram search for “pictorial balance” gives a good idea of this. Including only books scanned by Google, it exemplifies the above (see Figure 1). A theoretical explanation of how physiological interactions of visual data streams competing for attention create the illusion of there being a force field in pictures is described below

**Figure 1.**
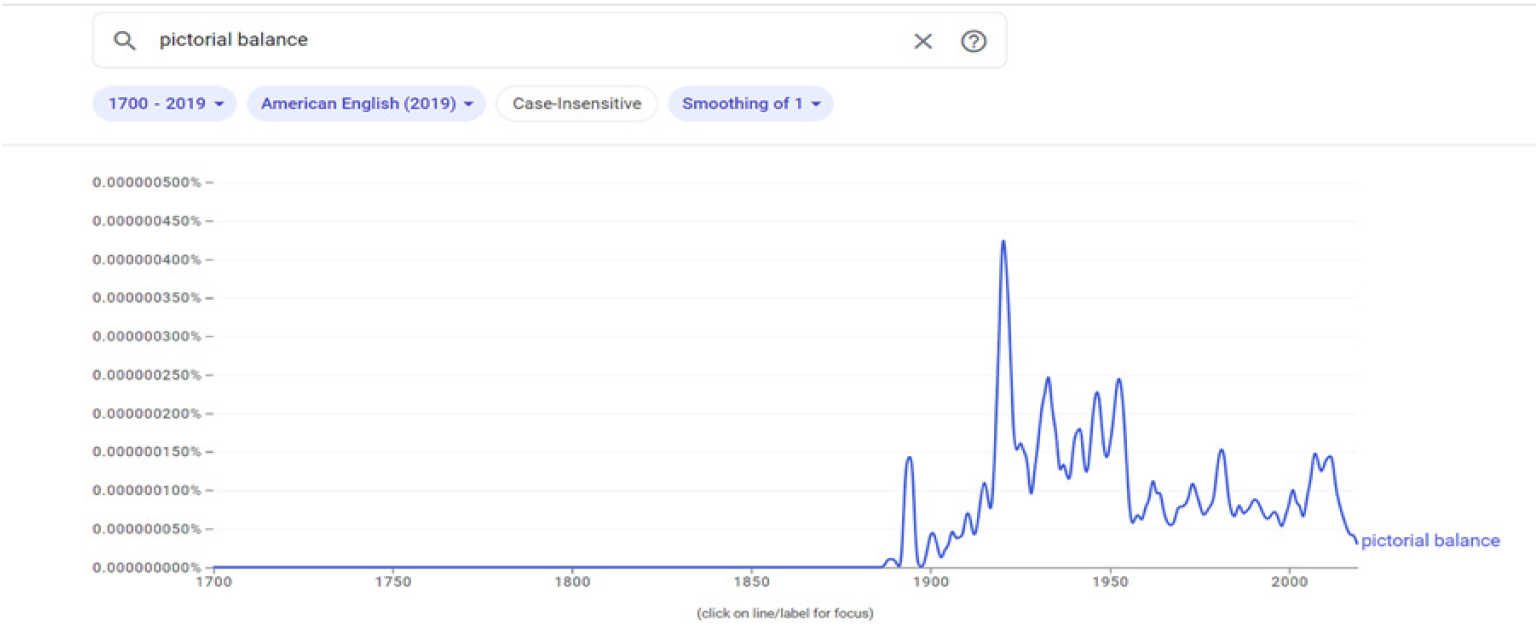
The frequency per year that “pictorial balance” was mentioned in books as scanned by Google

Balance has been thought to direct the eye, but it is salience that strongly influences eye movements. The initial sequence of saccades on first viewing a picture can be predicted and correlated with eye tracking through calculations based on the most salient features of surrounding areas ^12–14^. But for many years there have been studies that tried to show the more subtle effect of balance on eye movements. Langford could find no difference in these movements between pictures identified as either balanced or unbalanced. Locher and Nodine compared them in pictures that were thought to be balanced with variations of these pictures that were presumed less well balanced. They found that trained observers had more diverse and fewer specific exploratory movements in the more balanced compositions than untrained observers. On the other hand, in his extensive study of eye tracking, Buswell could find no difference between the two groups. The duration of fixations and the length of saccades are usually discussed in top-down cognitive terms of subject comprehension or artistic judgment unrelated to balance ^15–19^. As will be discussed the effect of balance is subtle and global in nature affecting ease of movement within the picture.

While this hypothesis of balance has been an intellectual interpretation of visual phenomena, the feeling of some sort of balance of visual forces requires an explanation of pictorial balance. To understand and measure low level balance, one must start with a picture that exhibits the sensation of perfect balance. Balance has been used to explain the feelings of pictorial unity and harmony, and there is an obscure effect observed by painters that does just that. There is no name or metaphorical description for the percept evoked by these pictures other than to say that they exhibit the aforementioned unnamed effect. It is usually only discussed in art schools when standing before such a picture and then in rather vague terms. The unexpected change in a painting from being remarkably luminously unified and harmonious to something less so or the reverse is quite striking. There is the feeling that the whole picture can be seen at one time and that the gaze moves smoothly and seemingly effortlessly without the apparent need to fixate on any pictorial element. There is a qualitative difference between perfect balanced where the gaze is looking at the picture as an object and an unbalanced painting where we are looking in the picture at the depicted forms.

Because there has been no name for this effect, historical discussion has been limited. Roger de Piles, an early 18^th^ century French critic and painter first described an arrangement of “lights and darks” creating an effect that is spontaneous and intense, riveting the attention and giving the impression that the viewer could gaze at the entire painting without focusing on any particular form. He thought this was the primary aesthetic effect of a painting ^11^. Similarly, Eugene Delacroix in the 19^th^ century wrote that “there is a kind of emotion which is entirely particular to painting: nothing [in a work of literature] gives an idea of it. There is an impression which results from the arrangement of colours, of lights, of shadows, etc. This is what one might call the music of the picture…you find yourself at too great a distance from it to know what it represents; and you are often caught by this magic accord ^11,20^.” Wassily Kandinsky in the 20^th^ century describes an analogous experience: “It was the hour of approaching dusk. As I was returning home … when suddenly I saw an indescribably beautiful picture, imbibed by an inner glow. First I hesitated, then I quickly approached this mysterious picture, on which I saw nothing but shapes and colors, and the contents of which I could not understand. I immediately found the key to the puzzle: it was a picture painted by me, leaning against the wall, standing on its side. The next day, when there was daylight, I tried to get yesterday’s impression of the painting. However, I only succeeded half-ways: even on its side, I constantly recognized the objects and the fine finish of dusk was lacking ^21 p.68^.”A contemporary description is provided by Brock: “some of [the photographers’] best work seemed to radiate a combined sense of balance, order, harmony and energy. The viewer’s eye seems to be held in the image in an almost hypnotic way, dancing around the picture ^22^.”

To study this effect, I created computer models that evoke it using experience from many years of painting. Image analysis indicated that it was perceived when a picture, centered at eye level and parallel to the plane of vision, has bilateral quadrant luminance symmetry with the lower half being slightly more luminous by a factor of 1.07 ± ∼0.03. A perfectly balanced painting will be described as coherent; all others will be called unbalanced for the study’s first objective. With respect to balance it was discovered by accident that the visual system cannot distinguish a picture from its white frame. I was printing black and white coherent pictures on white paper, and one time due to an error of alignment the picture appeared unbalanced. Due to the relative size of the picture and the paper I realized that the image would appear coherent whether framed or unframed unless there was this error of alignment. It was derived from this that with respect to balance, pictures are defined only by luminosity and not borders. For example, this is not true for an LED picture framed in black on a white ground where black marks the end of the picture.

There is a theoretical formula for balance which explains observations of a center-of-mass like effect using a modified variation of a standard center-of-mass calculation: if a rectangular picture with bilateral quadrant luminance symmetry and a darker upper half is said to be perfectly balanced around the geometric center, then the formula for the center of mass of four connected objects can be used. In this formula quadrant luminance L_xxQ_ replaces the mass of each object and is located at the geometric center of its respective quadrant. For simplicity, the picture’s geometric center is located at (0,0). L_TOTAL_ is taken as the average total luminance.

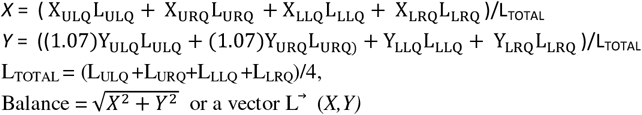

Y values of the upper quadrants are modified by 1.07 to account for the increased visual weight of the upper half. X_XXQ_ and Y_XXQ_ are the coordinates of the center of their respective quadrants, and L_xxQ_ is a quadrant’s luminance. The equation can be expressed as an addition of four vectors. Although this equation is precise for defining perfect balance, it is not clear whether it is true for states of imbalance. This will be called the geometric method of determining balance.

There is at least another way of calculating balance derived from the idea that it could be an early method for identifying and following moving organisms based solely on luminance. Since the start of this study I realized that although pictures are relatively recent, balance must have had a much older evolutionary significance. The luminance or visual method was suggested to me by the need to explain how the eye could divide a picture into precise quadrants by pure luminance. If a moving organism is defined as a collection of luminous points that move together with respect to a ground, it can be divided into two equally luminous parts by a virtual vertical line. The ability of the visual system to create virtual lines has been shown through the kanizsa illusion, and insects and evolutionary early vertebrates like sharks have been shown to see this^23,24^. Virtual parallel and perpendicular lines are drawn around the organism to create a “picture” while other diagonal lines determine the geometric center. This enables the drawing of a horizontal line bisecting the rectangle that divides it into four quadrants. The luminance of the upper half would be arbitrarily decreased by a factor possibly to account for light coming from above so as to render the organism more balanced, but more likely the opposite objective is required, so that the movements of an object appearing to have uniform luminance could be evaluated. The light energy or amount of light within a quadrant is then averaged over the area of the quadrant and treated as if it located at the geometric center of the respective quadrant or at the exterior corner of each quadrant. This permits calculations using the formula of the geometric method. Since there are two vertical lines dividing the organism, one into geometric halves and the other into two equally luminous parts, this algorithm provides two ways to calculate balance: the geometric and visual methods.

There are two effects conveyed by perfect balance: the intangible effort for the eye to move across the picture and the feeling that one can see the picture as a whole, riveting the attention. Both contribute to feelings of unity and harmony. When seen by reflected light, the two seem inseparable. However, it was observed that an LED picture is a poor conveyer of the attention-riveting effect while maintaining the effect on eye movements. This explains the study of Bertamini et al. who noted that observers preferred a print reproduction to the same picture as seen in a mirror or on a monitor and that they preferred the mirror view of the actual image to the monitor image ^25^. Locher et al. made similar observations^26^. The only difference between these three situations is that the light emitted from an LED screen is polarized. This is accounted for by Misson and Anderson who showed that polarized light increases luminance perception in the fovea and parafovea through unique macula pigmented structures that transmit polarized light and thus increase the sensitivity of central vision relative to peripheral vision ^27^. As a result of the altering effect of polarization, observer preference could not be used to show that the effect was seen in LEDs. However feelings evoked by the ease of movement in a coherent picture remain. This allows a performance based study to show that the effect was perceived in that two identical pictures differing only by balance could be seen as different. LED images had to be used to permit precise luminance determination and reproducibility under the conditions that were feasible. Perfect balance or pictorial coherence are used as absolute terms whereas balanced or unbalanced will be relative terms

The first object of this study was to show that some people are able to see this state of pictorial coherence by showing that they perceive a difference in two identical pictures differing only in that one is perfectly balanced. The second later objective was to show that one or both calculations of pictorial balance are valid by showing that the study group’s ability to distinguish differences correlates with balance as calculated by these definitions.

## Materials and Methods

Since a white-framed picture is seen by this low-level visual system as an object in contrast to the top-down view which sees it as a picture in an uninteresting frame, the comparison of a perfectly balanced picture with the identical but unbalanced picture can be done by a slight change of frame. If an observer compares carefully the two images and is able to disregard the frame change, any perceived difference would be ascribed to the effect. It was presumed that pairs of unbalanced pictures would be indistinguishable because I did not think that observers could see degrees of relative imbalance within the pairs. Many observers were excluded from the study because they did not pay full attention to the pairs. They would glance at the pictures rapidly and decide that there were no differences. Observers were included in the analysis if they carefully inspected the pictures, whether they saw any difference or not. Painters and those interested in looking at pictures were found to do this careful inspection. Others had trouble even when specifically asked although this was not usually done. Two studies were conducted using different frames. Each study consisted of the same ten pairs of images: five in which one of the images was perfectly balanced (balanced pairs), and five in which both were unbalanced (unbalanced pairs). Images were selected for their predominantly neutral content. They were prepared using Photoshop CS2^©^ in the following manner: quadrant image luminance can be changed in an unobtrusive and precise manner using the lasso tool to create an irregular form within a quadrant and changing the luminance with the level command by no more than ±5 percent. This form is then moved around and the operation repeated until the desired quadrant luminance value is reached. The two studies differed only in their use of different sets of white borders. The three possible image and border combinations are in Figure. 2. Study I compared Figure 2a with either 2b or 2c. Study 2 compared Figures 2b with 2c. The observer viewed sequentially a picture with different frames on a color-calibrated iPad3 using the ColorTrue™ app (which is no longer available) and a Colorite™ calibration device. The observer viewed the iPad centered and parallel to the plane of vision at arm’s length. The images on the iPad were approximately 5 × 6.75 inches. The pictures used for the studies are shown in Fig. 3, and this is the order in which they were seen. Images 1,3,5,7, 8 were the balanced pairs. (all images are in the supplemental files labeled with quadrant luminance data).

**Fig. 2.**
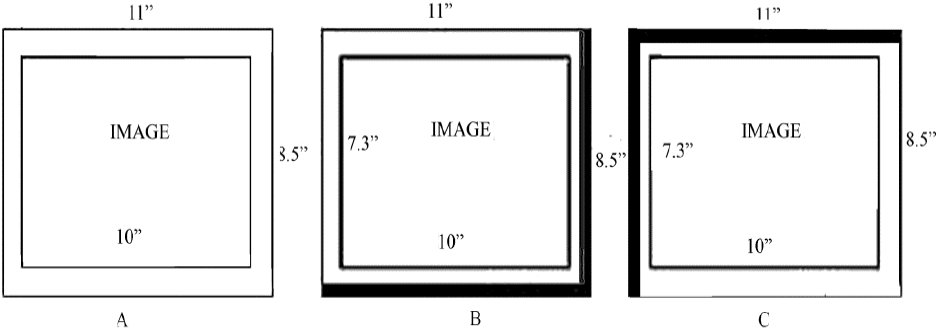
The three possible image and border combinations: the wide black lines would not be seen on a black ground.

**Figure 3.**
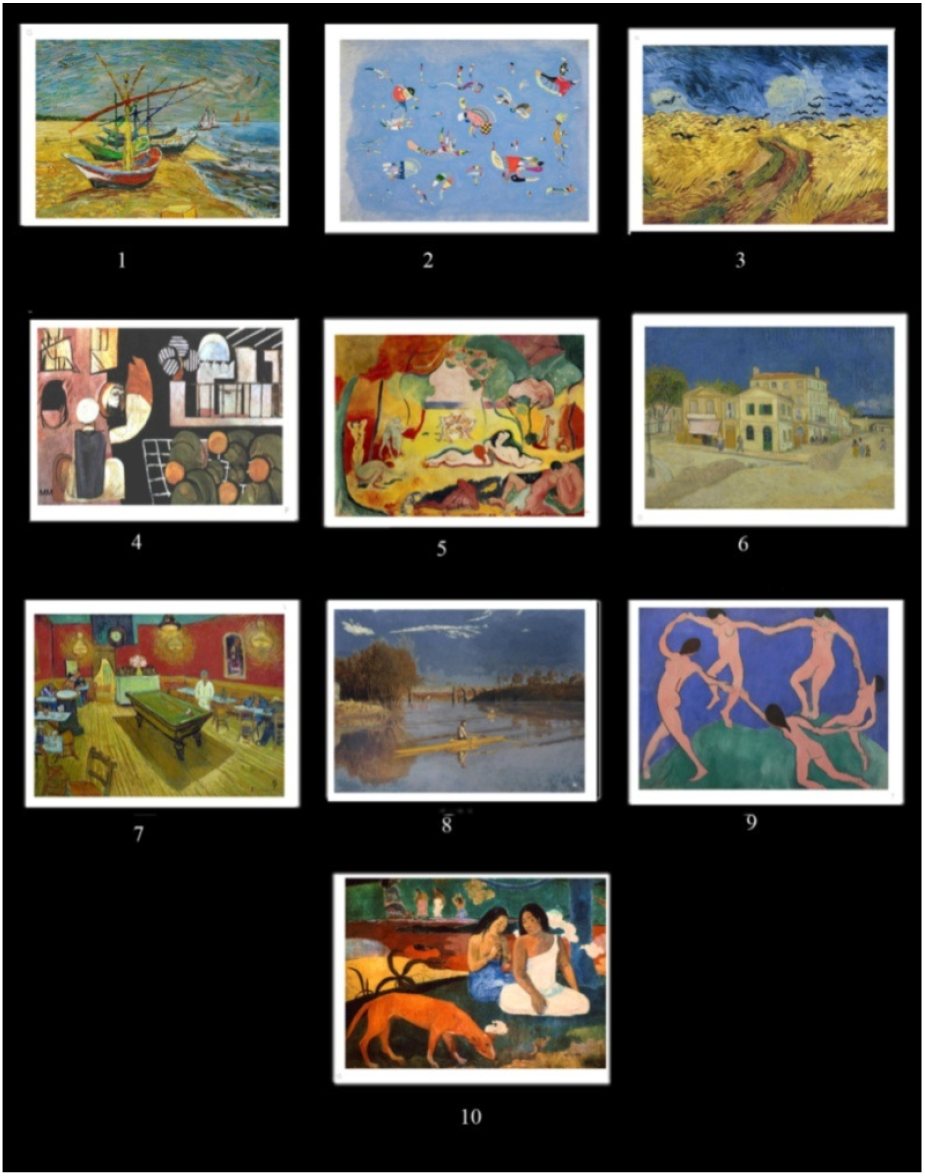
Pictures used in the study pairs

The observers gave their verbal consent to participate anonymously in a study about how people look at pictures. They were told that it would consist of looking carefully at ten pairs of pictures and being asked whether the central images appeared to be the same or different while disregarding the frames. They were also told that it was intended for some pairs to be seen as the same while others as different. No identifying information or medical history was obtained, and there was no attempt to eliminate observers who were color deficient, had cataracts or impaired binocular vision. 45 observers were included in the first study and 39 observers in the second study. Observers were excluded if they did not follow the directions by either rapidly saying without careful examination that they could see no difference or by insisting that the frame change was the only difference. There were only four of the latter. For the unbalanced pairs it was presumed that because of only slight differences of imbalance observers would find the pictures to be identical so that the response of “same” was considered correct. With the balanced pairs a response of “different” would be labeled correct indicating they had seen the effect. Observers viewed the pictures sequentially on an iPad at arm’s length while I sat slightly behind them. They were permitted, indeed encouraged, to hold the iPad themselves as long as it was maintained correctly centered and parallel to the plane of vision. There were no restrictions on the length of observation, and they could return to a previous image as much as they wanted. However, subjects were told that the difference if any was more in the way of a feeling and were discouraged from making point by point comparisons. The difference was described as analogous to that between monophonic and stereophonic music: same music but seems different.

## Results

Four observers could identify all the balanced pairs correctly, and 5 observers made one error (Table 1). Some subjects thought they saw differences in color while one thought the depth of field was different, and many felt there were differences but could not describe them. A simple analysis of the study could not prove that observers perceived the percept of perfect balance. However, a second analysis including the calculations of balance (table 2) shows that either the observers saw the effect of perfect balance and/or the effect of relative balance. Table 1 shows whether they could see intra-pair differences while table 2 shows that they can see inter-pair difference

**Table 1.** the number of observers correctly answering the same/different question with the 10 pairs.

| 1st study |  | 2nd study |  |
| --- | --- | --- | --- |
| Observers | Correct Responses | Observers | Correct Responses |
| 10 | 3 | 10 | 1 |
| 9 | 2 | 9 | 3 |
| 8 | 2 | 8 | 1 |
| 7 | 4 | 7 | 6 |
| 6 | 13 | 6 | 9 |
| 5 | 12 | 5 | 11 |
| 4 | 7 | 4 | 5 |
| 3 | 1 | 3 | 1 |
| 2 | 1 | 2 | 2 |
| Total | 45 |  | 39 |

**Table 2.**
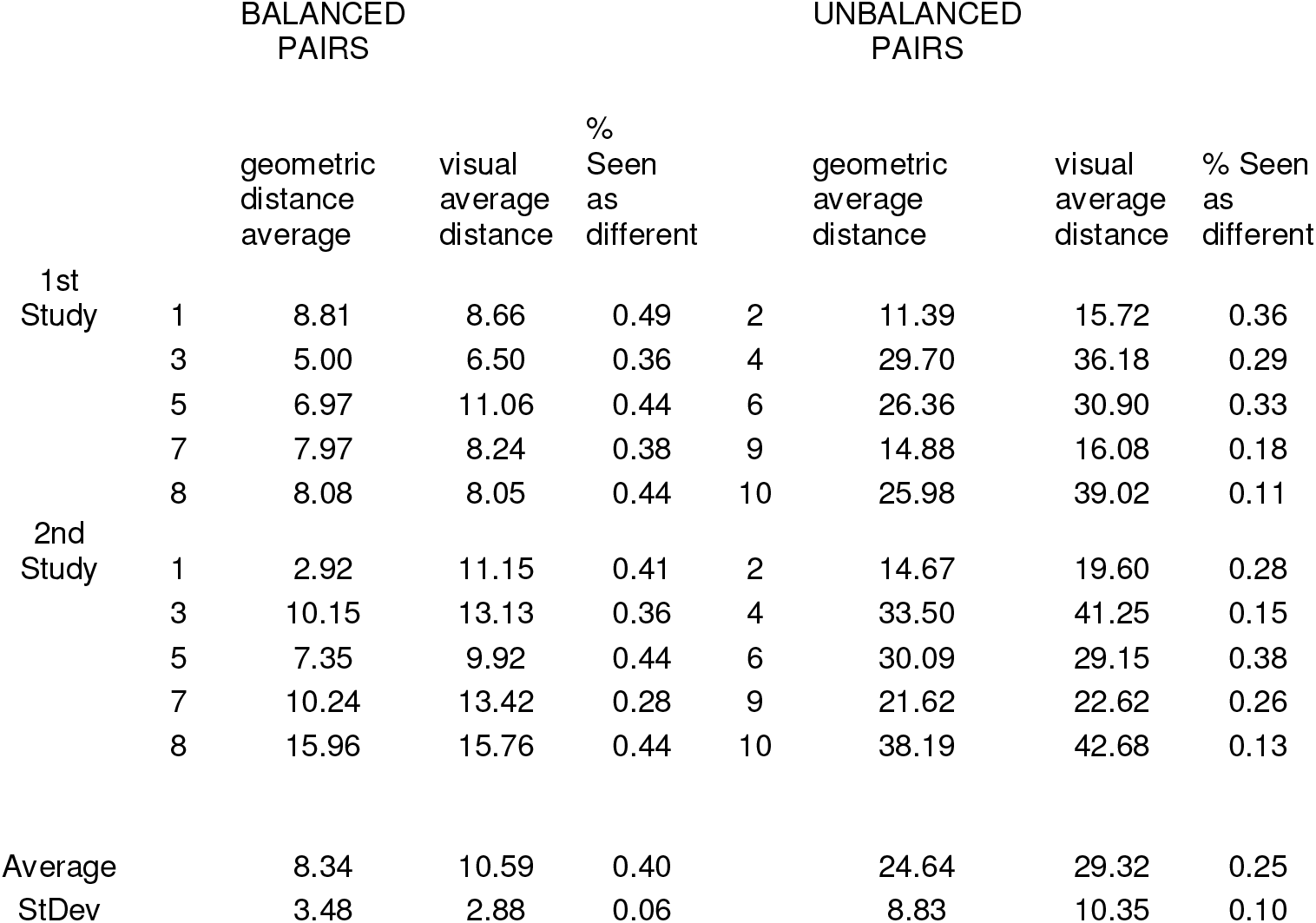
Pair average distance and percent of observers who identified the pairs as different using both methods of balance. The MATLAB™ code used to determine visual balance and files of the corresponding pictures without black borders are in the supplemental files.

It was noticed to my surprise that subjects saw differences in the unbalanced pairs. At first I thought this was due to guessing, but on further examination it was found that there is an inverse correlation between the percent of pairs seen as different and the pair’s average distance (average imbalance) as calculated with the two methods (see table 2). This can be appreciated if the pairs are divided into groups based on their degree of average balance where there is clearly a group in which the average state of imbalance is small (the balanced pairs), and the another where the average imbalance is large.

The correlation for the geometric determination between average pair imbalance and the percent identified as different is: r (18) = −0.676, p = 0.001. The same correlation for the visual determination is r(18) = −0.757, p <0.001. Figure 3 shows the results graphically.

**Figure 3.**
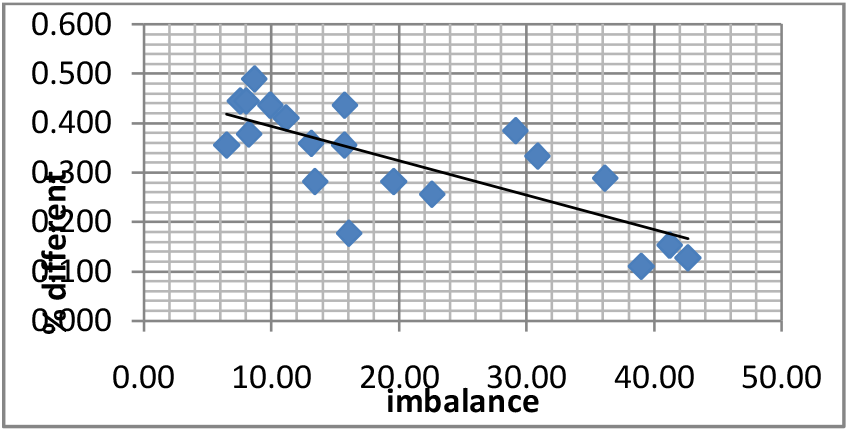
Graph of the visual balance results

These finding suggest one or both of two interpretations: either the observers are seeing the effect of coherence to distinguish the pictures within the pairs demonstrating that they see the effect, or the observers are seeing pairs according to their average state of balance. The second point is reinforced by the following: if the average of the balanced pairs is used as one data point so as to avoid the statistical influence of their relatively large number and the ten unbalanced pairs as other data points, the visual method has a correlation of r (11) = −0.64 p = 0.046. Thus subjects most likely perceived relative average balance across the pictures and not just between pairs with low and high average balance. This correlation of pair average balance with observations of same or difference is not obviously logical. As I wrote above, I thought that observers would be comparing intra-pair difference in which two unbalanced pictures would seem identical, and I only found a correlation between inter-pair average luminance and observations, Why should the results correspond to different degrees of viewing pairs as different based on average states of balance? An ecological interpretation of the study discussed below explains this.

The results are true for the observers selected for their ability to follow the rules who were mostly painters. This correlates with studies showing that painters view pictures differently than untrained individuals ^28–33.^ Antes showed that fixation patterns distinguish painters from non-painters, and Koide et al. using a prediction model of a saliency map showed that artists are less guided by local saliency than non-artists ^34,35^. It was noted that painters become much more involved in the act of looking than non-painters, and that subjects frequently complained that it was tiring to do the study. It is hard work to look carefully at pictures. Moving the eyes seems effortless, but within pictures it is different because of balance. The reason for this will be discussed below.

Although a few non-painters were found to discriminate between the pictures, any study is limited by the need to find large numbers of painters. The use of an iPad in different lighting environments was necessary to bring the study to the observers as no laboratory was available. Reflections from the iPad screen increased the number of false negatives, i.e. coherent pictures being seen as unbalanced as the reflections change quadrant luminance. It is highly unlikely that the reverse would create states of balance that correspond to the results because observers did the study in many different areas. That observers saw unbalanced pairs as being different was contrary to expectations, and therefore could not have been influenced by the examiner. The methods for calculating balance were determined after the study was done. It has been suggested that they were created so as to determine the desired results disregarding the logic of the formula and algorithm, but one would have to explain how this was done as they gave paradoxical results..

## Observations

The characteristics that I have observed of a picture evoking the percept of pictorial coherence indicate that it is extremely sensitive to small differences of quadrant luminance. This explains its fugacity without precise viewing conditions. I have observed that with pictures in a studio the change in luminosity of drying paint several minutes to hours following the creation of a picture is sufficient to to cause the effect to appear or disappear. Changing the illuminating spectrum or the painting’s position relative to the viewer destroys the effect. Given that many do not see the effect, and that it is normally created through chance and even then is fleeting and that there is no name for it, one can understand why it is not discussed in 20^th^ century literature. Although the formula for coherence has been defined for viewing a picture directly in front of it at eye level, a given picture can be seen as coherent from another set of conditions. It is possible that given the proper frame many pictures might be seen as coherent when directly in front of the viewer.

Pictures with many equally prominent forms have the same or similar effect as poor balance. Pair 2 is a good example of this. The eye is forced to go from form to form with perhaps slightly greater ease as balance improves. However, if the picture is seen from a distance, the forms coalesce so that the picture appears coherent. Prominent geometric forms enclosed by definite lines and flat color will prevent the painting from appearing coherent. However, if the circumscribing line is imprecise and the interior surface is variegated, they may be part of a coherent painting. For example, the paintings of Mark Rothko often present rectangles laid on a ground. However, in general, all the borders are imprecise and the surfaces are variegated so that those of his paintings keeping closely to this style can be seen as coherent. An analogous situation is that of a flat surface with no marks, i.e. one coated with an opaque layer of paint gradually modified so that the upper half is slightly darker. If one calculates backwards from the perceived balance i.e. the originally viewed object unmodified by balance calculations, it would be a uniformly luminous object. There has to be some surface quality for an object to be balanced. It has been shown that forms in nature have a fractal quality, that visual systems have evolved to respond to conditions of nature that are fractal, and that fractal images have an aesthetic quality ^35–37^. Therefore, it can be inferred that the part of the visual system that calculates balance is also most sensitive to fractal forms. The visual system is designed to work best in this fractal environment and not the world of manmade objects that are linear and uniform, causing the eye to jump from one form to another. In addition, it has been noted that binocular vision is necessary to view the percept.

The physical evolution of eyes of organisms in a water environment has been described in considerable detail starting from light sensitive cells to organs that map the light from the exterior to a light sensitive retina. However, it has not been discussed how an organism can make sense of this luminous flux. Here it is proposed that this is done through the calculation of balance. A moving organism is identified by the calculation of a vector of balance derived from its virtual rectangle. Once the measurement of balance is done, the organism can be followed so that a complex object of whatever size is reduced to an ever changing vector and a rectangular halo. Even if it stops moving, the vector and halo would remain visible where it might otherwise blend in with the ground. Many different organisms can be followed simultaneously with minimal computing power, and it guarantees that at any moment either the prey or predator is located within its boundaries.

Evaluating an organism in terms of balance means that it is always viewed at any given time as if it were a two dimensional object directly in front of the viewer. A demonstration of this occurs in the phenomenon sometimes called the Mona Lisa effect that has never been adequately explained in which the gaze of a depicted person looking straight at the viewer positioned before the picture seems to follow them as they walk to the side. The painting, *Mona Lisa*, only imperfectly exhibits this effect, but there is no other name^38,39^. To view a picture from the side does create a distortion due to foreshortening such that it should appear to be a trapezoid with the interpupillary distance of Lisa appearing slightly diminished. However, the low level view fills in the spaces to create a rectangle so that this foreshortened picture with the virtual borders appears like a picture directly in front of us with Ms. Lisa staring at us as before (Figure 4).

**Figure 4.**
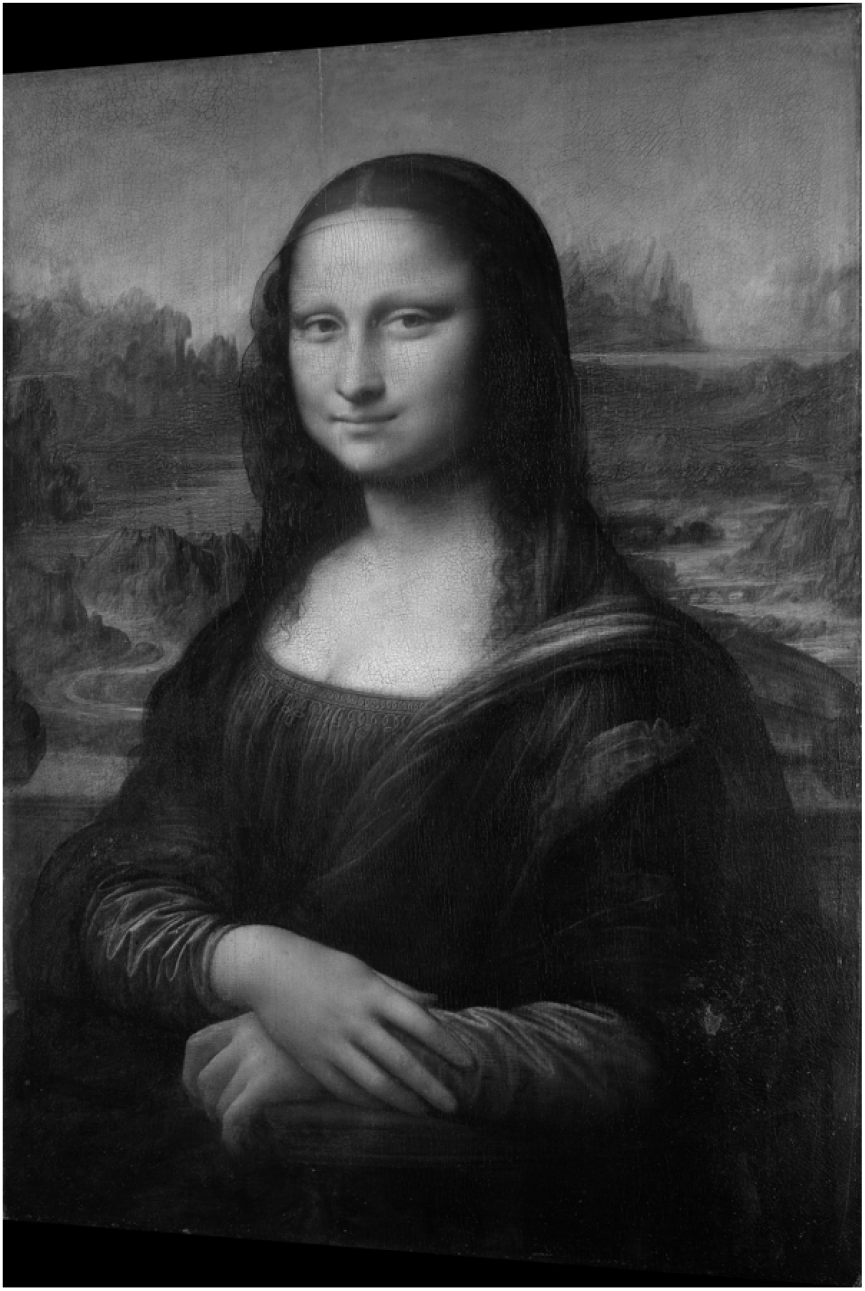
A simulation of *Mona Lisa* viewed from the side at eye level.

As previously noted the results of the study are not logically apparent. Why should average pair balance correlate with the degree of appearing different among the ten pairs (inter-pair difference) when the subjects are asked to evaluate intra-pair difference? An ecological explanation in which the pictures are themselves interpreted as organisms explains this. A bilaterally symmetric organism or its fairly balanced pictorial equivalent would be seen as possibly approaching so that it would require particular attention to be directed at the two different but similar images to evaluate what might seem threatening. However, as the pairs become unbalanced, they would be interpreted as the same organism moving somewhat orthogonally with no threatening or other implications.

I began this paper with a discussion of the 20th century concept of balance starting with Poore’s observation that “…every item of a picture has a certain positive power, as though each object were a magnet of a given potency ^5^.” Fifty years later Arnheim described similar perceptual forces of a picture that he attributed to their location in the picture ^2^. I also described the effort I experienced moving my eye around the picture without discussing pictorial forces. The reason for these sensations is the following: When viewing a picture, the information about balance from peripheral vision uses the tectopulvinar pathway that directly accesses the pulvinar, a central point controlling attention (this is looking at) ^40–43^. It is in competition for attention with the parvocellular stream from the fovea using the geniculostriatal pathway, and it suppresses information from the magnocellular stream which derives information from the same peripheral retinal cones. In a balanced picture, we are looking at the object using peripheral vision, and it appears as a luminous unified whole. In an unbalanced picture, we are looking in, and central vision dominates, suppressing peripheral vision. Since peripheral information is needed to facilitate saccades, the eye feels stuck ^44^. This increase in ease of movement in a balanced picture has been interpreted as the result of forces in equilibrium around the geometric center of the picture.

The attention-riveting effect of a perfectly balanced picture is an artistic peak effect, different but similar to the peak effects in other art forms which can be riveting in their own way ^45,46^. A “peak artistic experience” in this context is characterized by the work commanding total attention, and as such it can be a dissociative experience such as described by Brantley in his review of *Uncle Vanya* ^47^. Such an experience is always felt as an intense surprise, and a surprise can only be described using an analogy to an equivalent surprise. No one is so highly sensitive to every art form. Clive Bell in his book *Art* describes his marked aesthetic sensitivity to painting and compares this to his relative lack of sensitivity to music ^48^. He can enjoy and intellectually understand why a musical composition or performance is excellent, but it is not the same as his response to painting. He can make this distinction only because he has this acute sensitivity to painting and its aesthetic percept, to which he can compare his responses to other art forms. Without that he might have expected that any aesthetic response is an intellectual one. The distinction between a low level aesthetic percept and higher level intellectual aesthetic responses is fundamental to the understanding of art and a source of much confusion.

This paper reintroduces the percept of pictorial coherence as first described but not named by Roger de Piles to explain the feelings of unity and harmony evoked by certain pictures^11^. It proposes a calculation of pictorial balance using an algorithm that can also be interpreted as a method for a primitive organism to organize the raw perception of light to see and assess the movement of other living organisms. It further proposes a way for this primitive luminance visual stream to interact with evolutionarily later visual streams to enable us to perceive pictures as aesthetic objects. The interpretations of the study and its theoretical conclusions have an internal logic, but further experimental evidence will, of course, determine its validity and how it affects us in other aspects of vision. A new percept requires a modified theory of vision and introduces a different conception of the aesthetic experience. It should be an important neuropsychological objective in other genres to understand what and how an ordered collection of lights, sounds and movement can attract the attention so intensely on a pre-cognitive level in people as to induce an intense dissociative type of experience that they call an aesthetic experience. This might not only enable us to understand our experience of and judgments about a work of art but provide greater understanding of how the mind processes sensory experiences.

The Matlab code for calculating the visual method of balance was largely provided by the Matlab Mathworks Help community. This has been verified with a script from CHATgpt.

Supplemental Information is available for this paper including all documents and information necessary to repeat the study and confirm the results using a color calibrated monitor. Additional picture pairs will be provided on request.

