## Supplementary material for "Pictorial balance, a bottom-up neuro-aesthetic property mediating attention and eye movements, explains the feeling of unity and harmony in pictures. A primitive visual operating system determines balance and shows how the world was first visually organized using luminance and movement": MatLab script for Visual Balance.m

```
A = imread('picture.tif');
A = rgb2gray(A); % BT601 luma
A = im2double(A);
% split the image into two halves of approx equal weight
Aps = cumsum(sum(A,1),2); % cumulative sum of image profile
lastidx = find(Aps<=(max(Aps)/2),1,'last');
Aw = A(:,1:lastidx); % west half
Ae = A(:,lastidx+1:end); % east half
% show the weights are close to equal
[sum(Aw(:)) sum(Ae(:))];
% bisect halves
hh = round(size(A,1)/2);
Anw = Aw(1:hh,:);
Asw = Aw(hh+1:end,:);
Ane = Ae(1:hh,:);
Ase = Ae(hh+1:end,:);
% average luma for each quarter [nw; sw; ne; se]
allluma = [mean(Anw(:)); mean(Asw(:))
    mean(Ane(:)); mean(Ase(:))]
% find geometric center vectors
center = [lastidx hh]; % [x y]
allv = [center/2;			% [nw; sw; ne; se]
    lastidx/2 hh*1.5;
    lastidx*1.5 hh/2;
    center*1.5];
allv = bsxfun(@minus,allv,center).*[1 1.07; 1 1; 1 1.07; 1 1]
% weight center vectors by luma
allv = bsxfun(@times,allv,allluma)
sumv = sum(allv,1)
```
