## Supplementary figures and images for "Pictorial balance, a bottom-up neuro-aesthetic property mediating attention and eye movements, explains the feeling of unity and harmony in pictures. A primitive visual operating system determines balance and shows how the world was first visually organized using luminance and movement"

### 01 150.79 149.02 151.12 147.81 (586x783) lower.tif

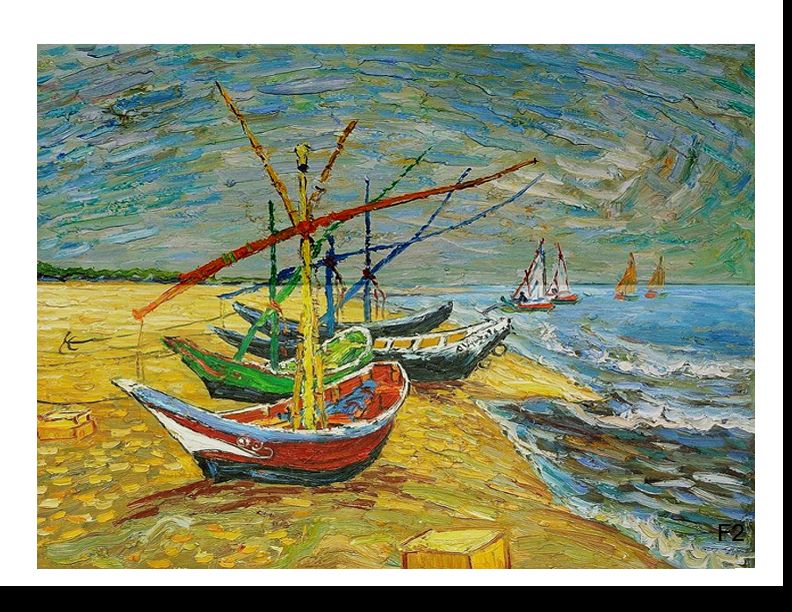

### 01 150.79 149.02 151.12 147.81 (586x783) lower.tif

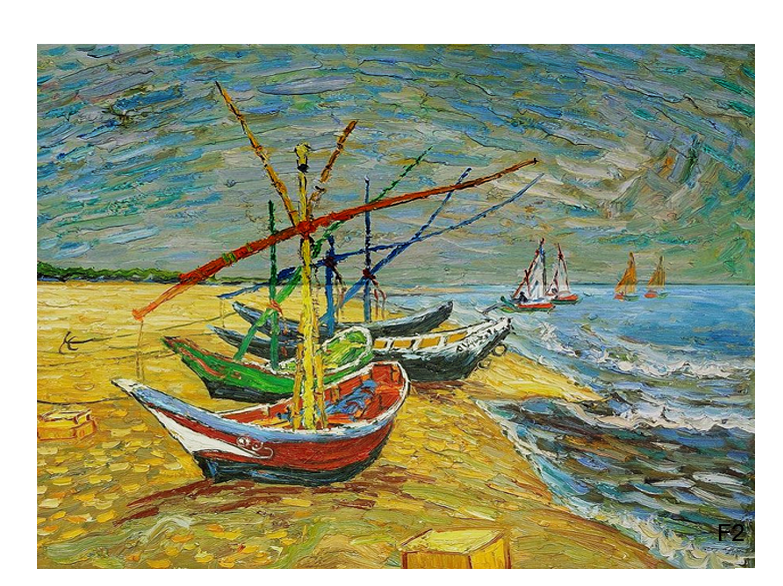

### 01 155.49 148.62 151.82 144..68.tif

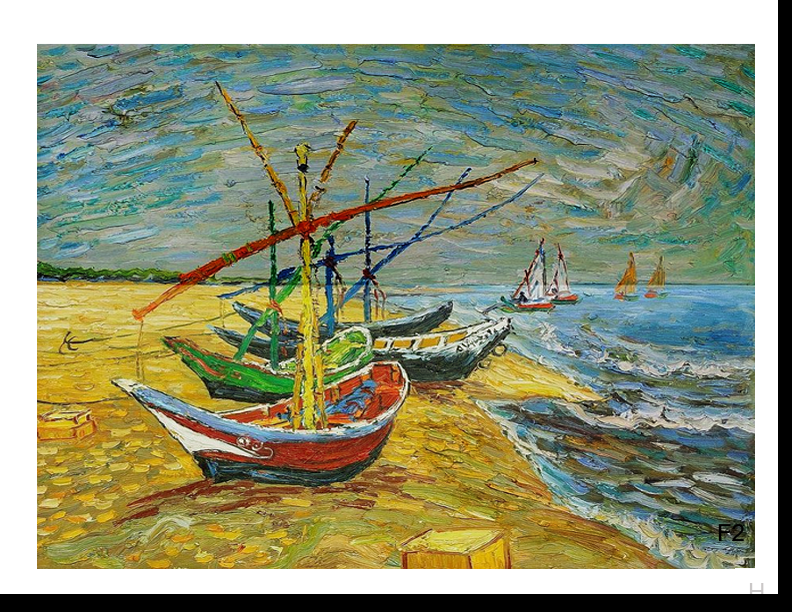

### 01 155.4477 148.5839 153.5604 144.5558.tif

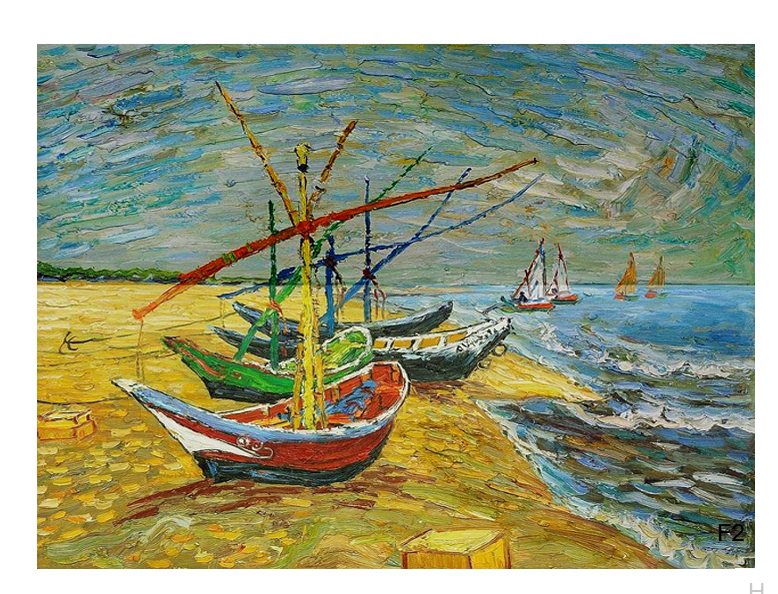

### 02 146.30 146.25 154.98 155.07 1.06.tif

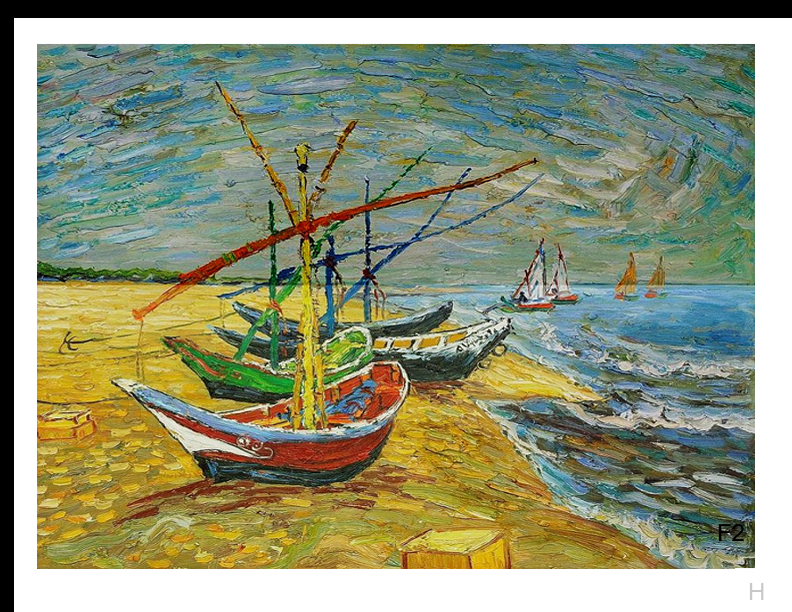

### 02 146.30 146.25 154.98 155.07 1.06.tif

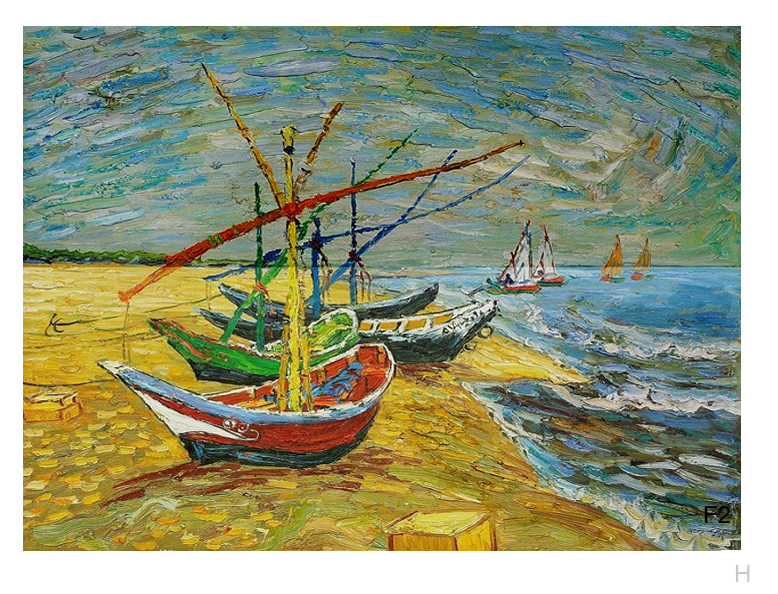

### 03 174.26 174.52 174.09 (586x783).tif

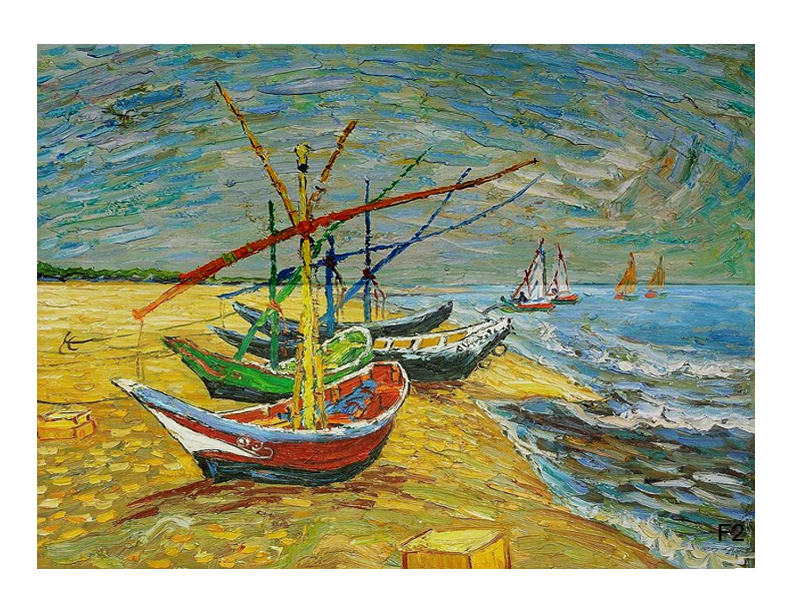

Supplement: Supplemental Files [file 104687_file03.zip › 20 Pictures used in 1st study including black bars/02 151.53 151.58 158.97 159.06 {[1.049].tif]

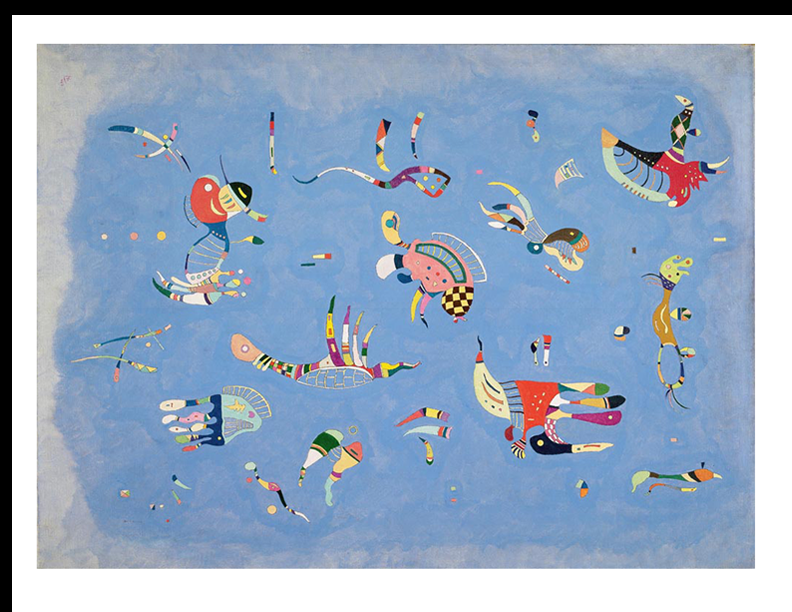

### 03 174.26 174.52 174.09 (586x783).tif

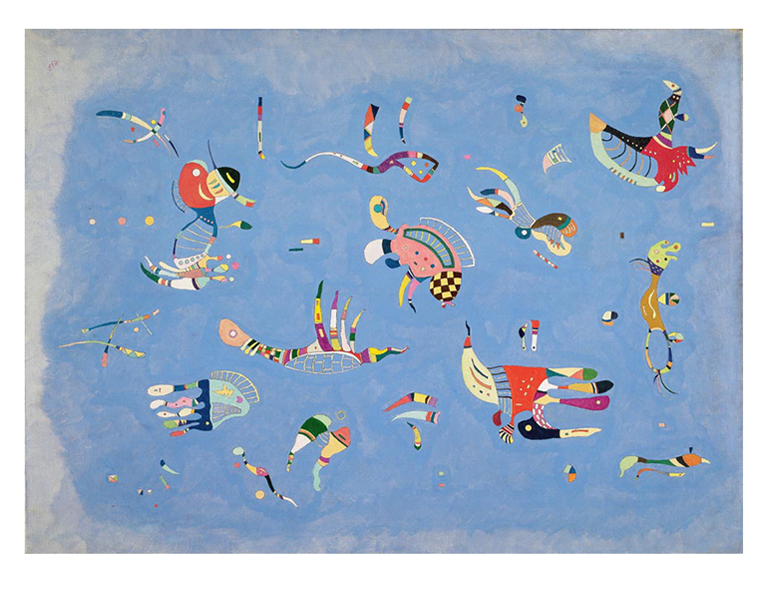

### 03 181.4 175.9 171.5 166.8.tif

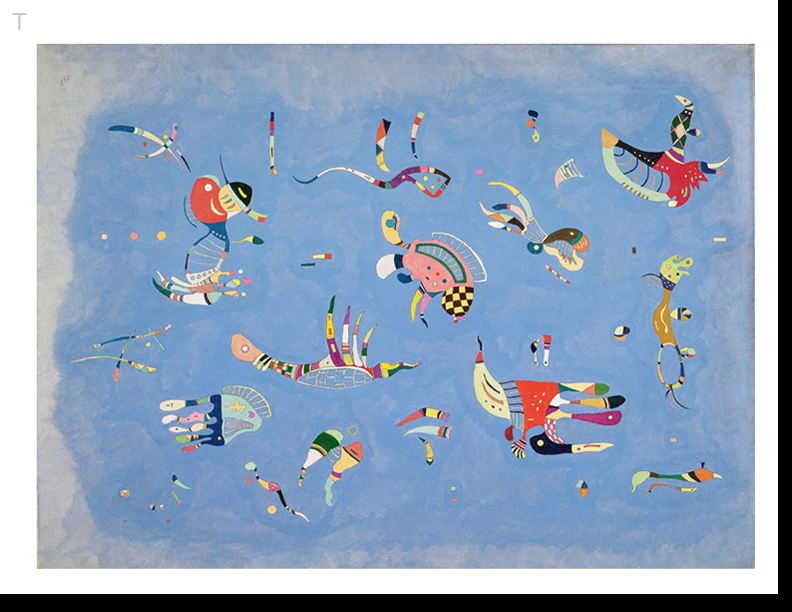

### 03 181.4 175.9 171.5 166.8.tif

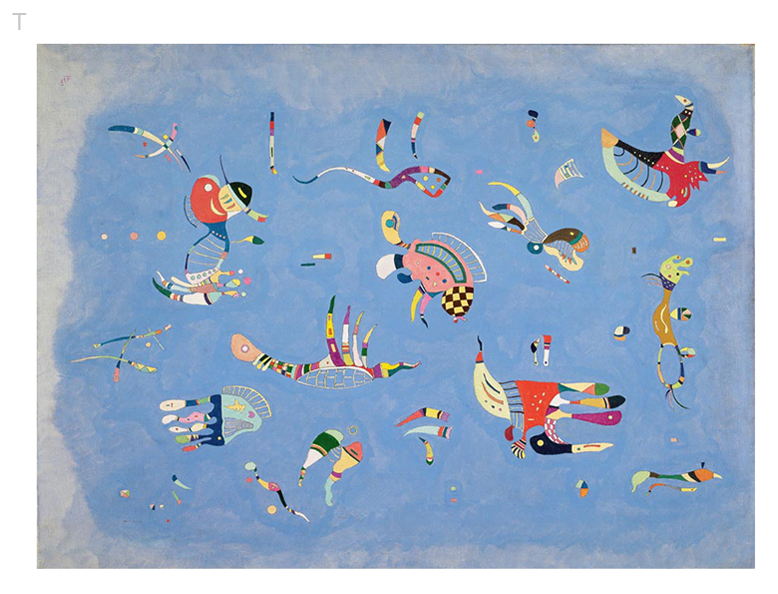

### 04 172.66 173.5 173.7 175.7.tif

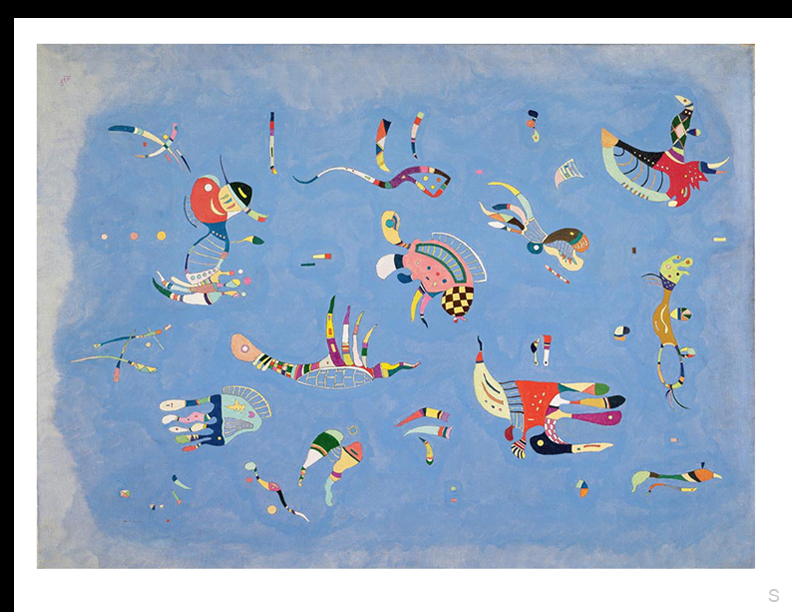

### 04 172.66 173.5 173.7 175.7.tif

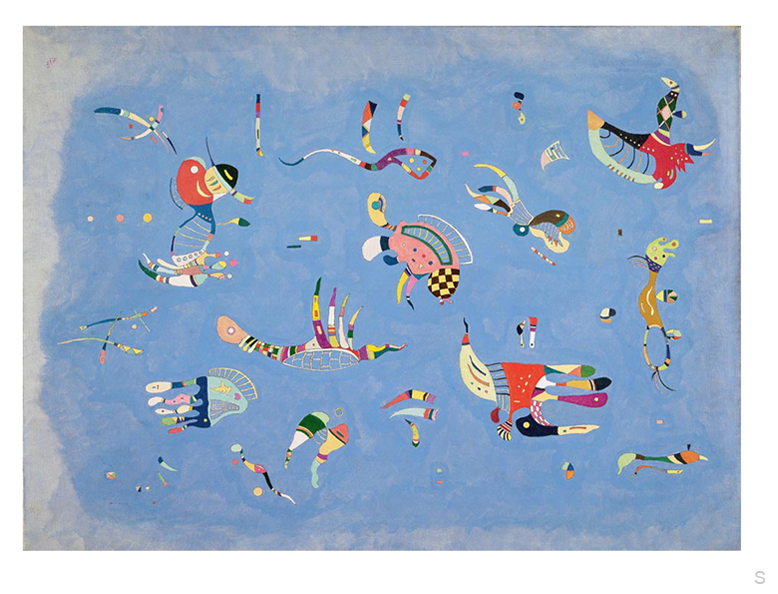

### 04 179.22 176.65 176.29 175.14.tif

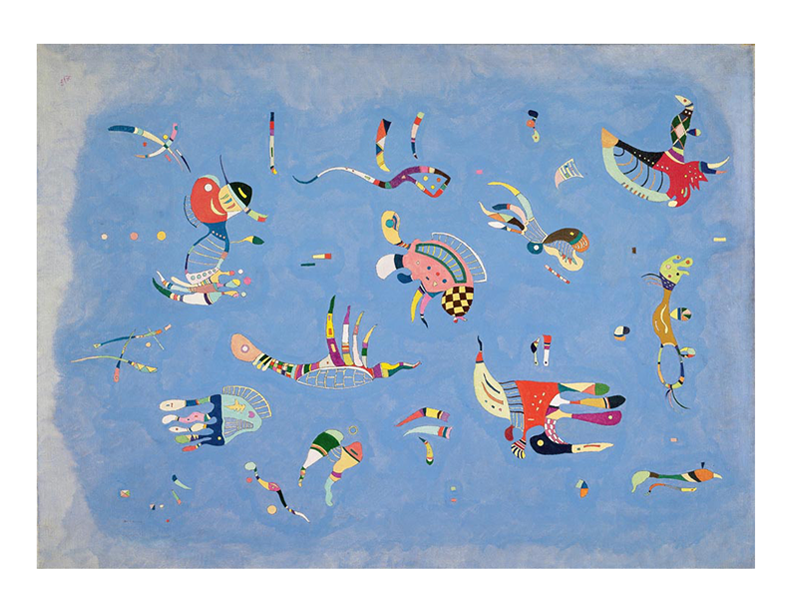

### 05 142.37 142.07 150.28 150.18 1.056.tif

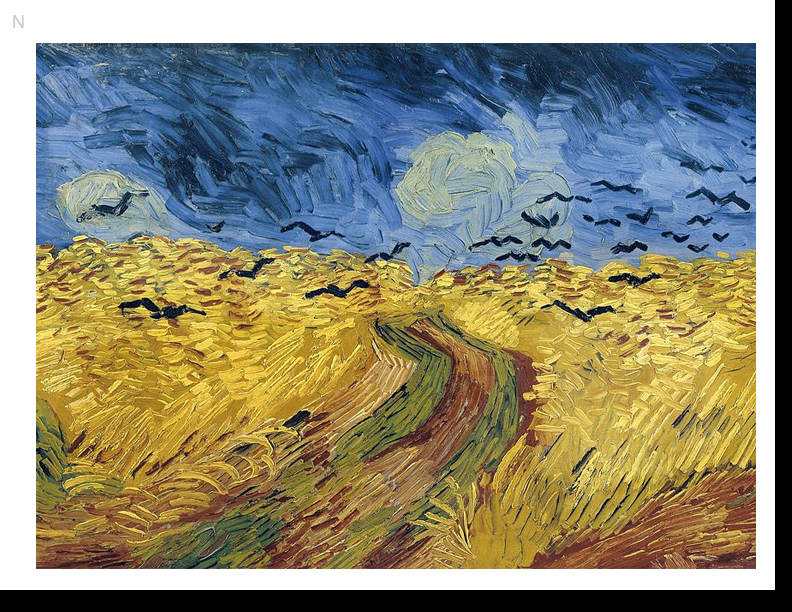

### 05 142.37 142.07 150.28 150.18 1.056.tif

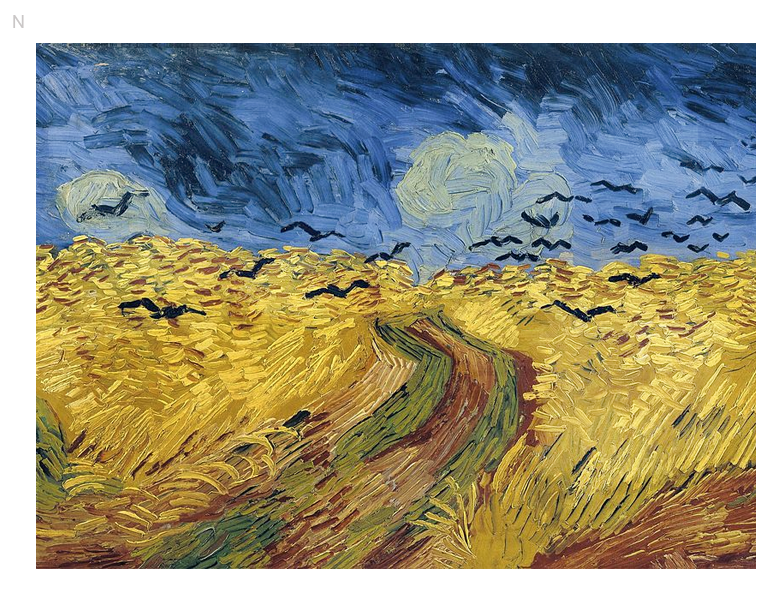

### 05 142.96 148.72 157.39 162.31.tif

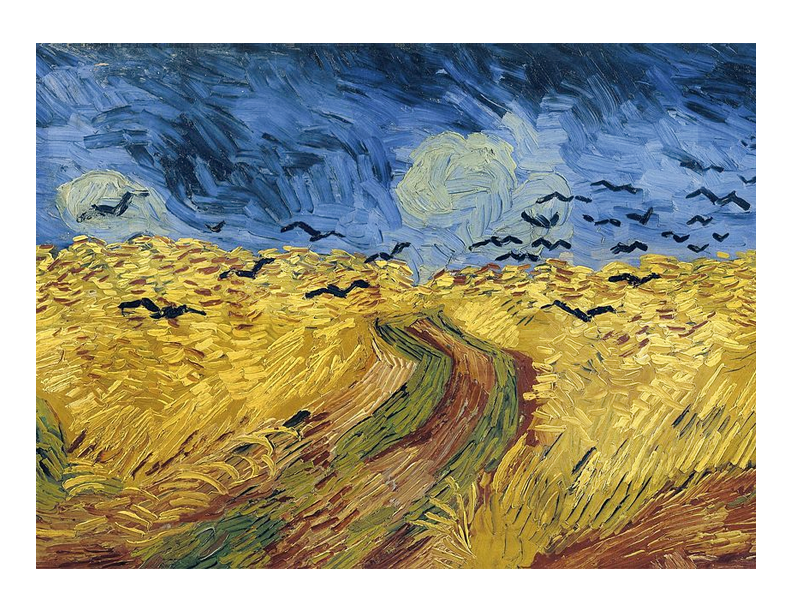

### 06 130.33 142.06 152.66 162.76.tif

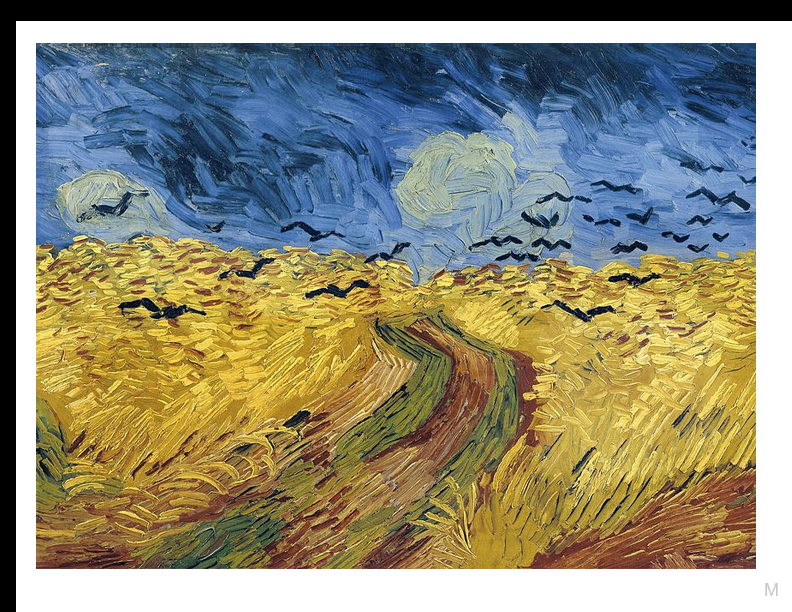

### 06 130.33 142.06 152.66 162.76.tif

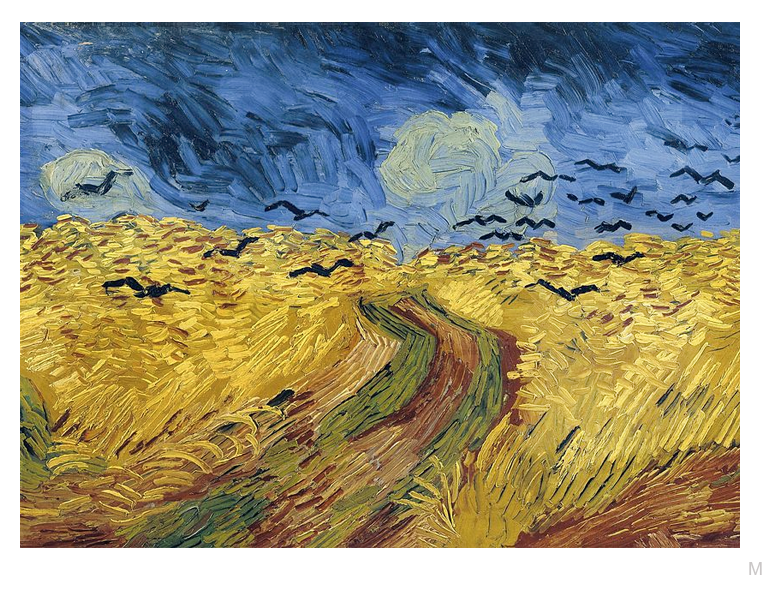

### 06 142.37 142.07 150.28 150.18 (590x775) 1.056.tif

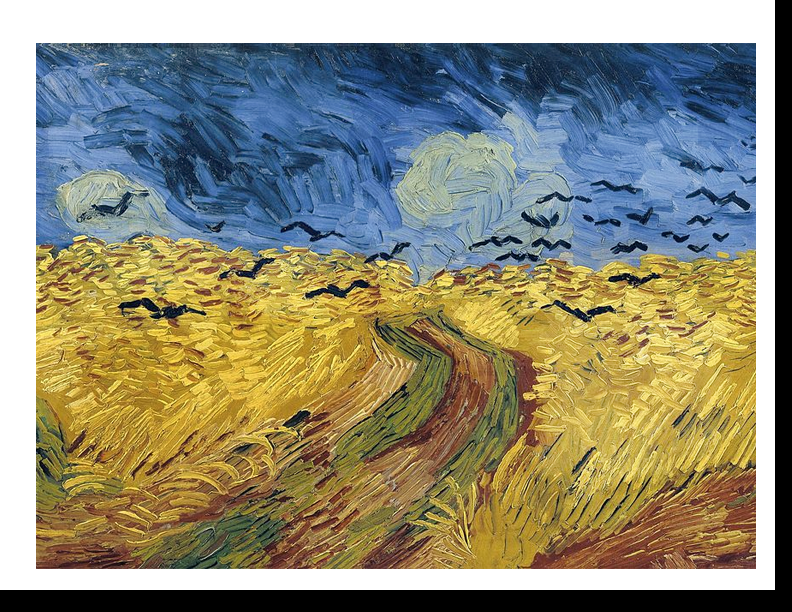

### 06 142.37 142.07 150.28 150.18 (590x775) 1.056.tif

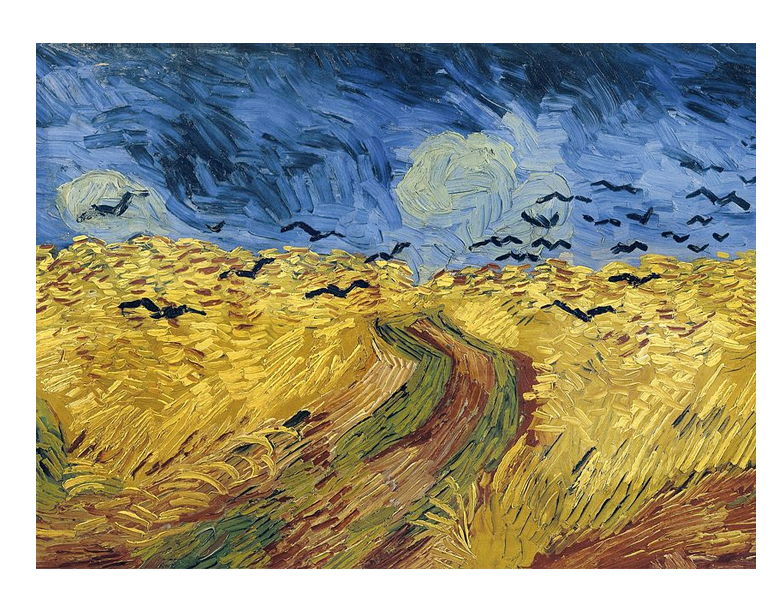

### 07 153.37 161.65 144.09 140.46.tif

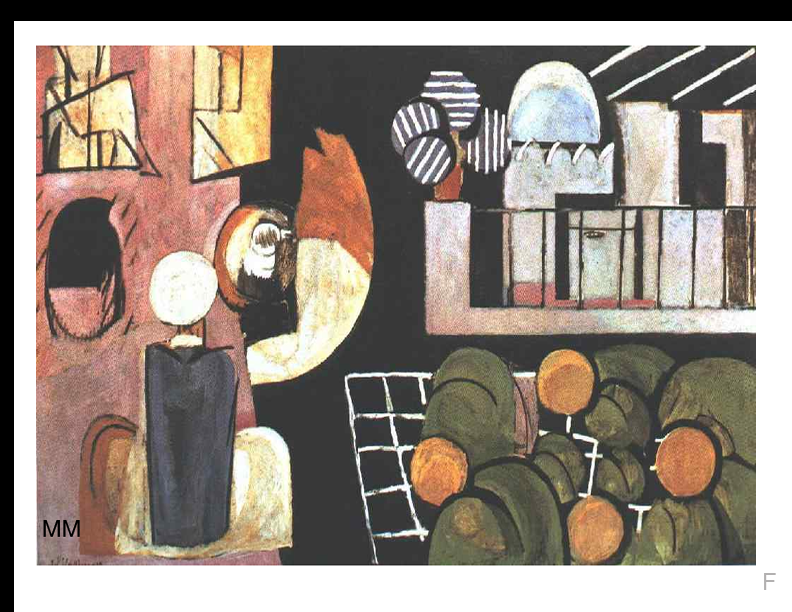

### 07 153.37 161.65 144.09 140.46.tif

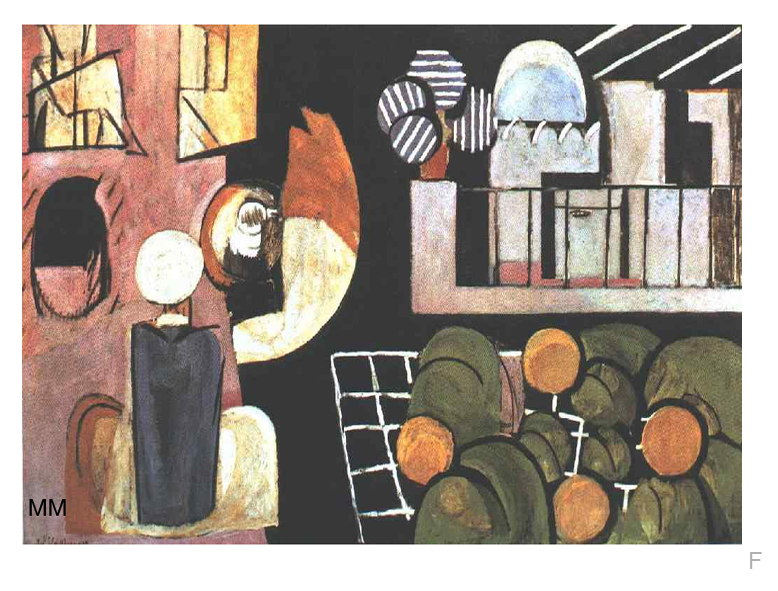

### 07 162.98 153.38 141.04 142.83 595x774 upper.tif

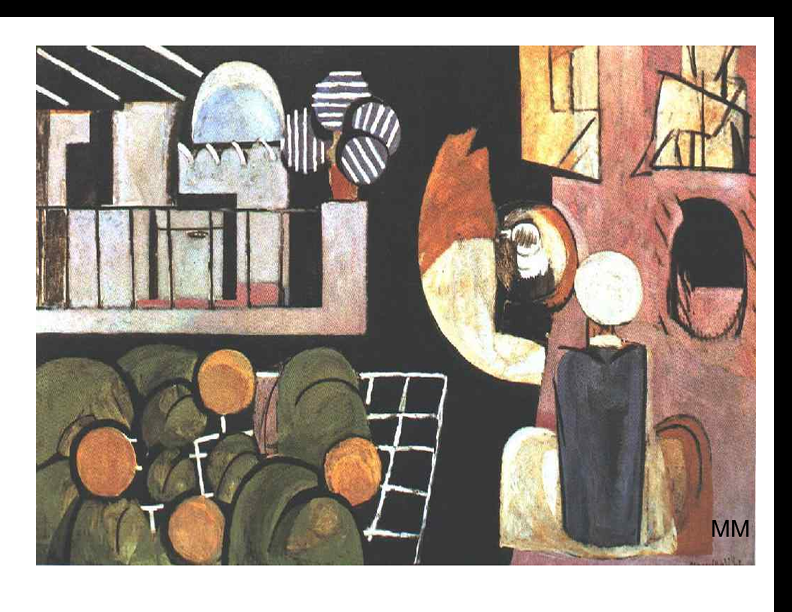

### 07 162.98 153.38 141.04 142.83 595x774 upper.tif

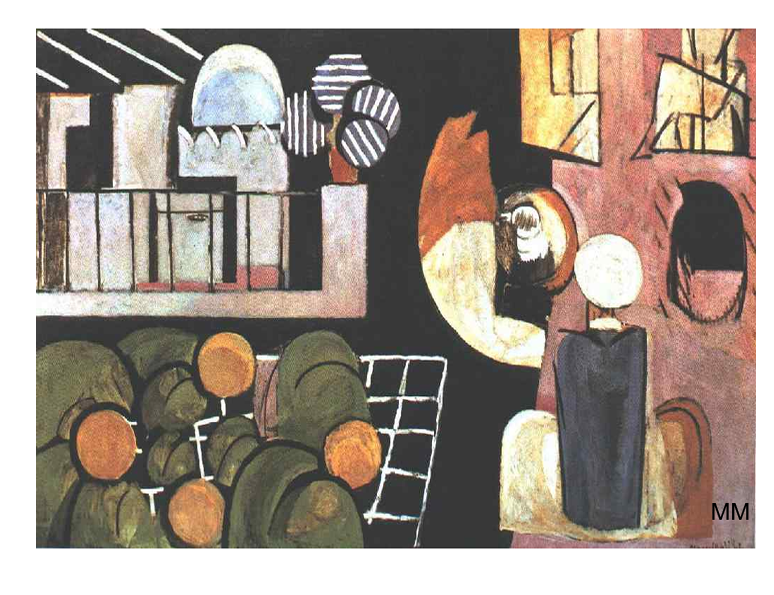

### 08 165.94 162.77 142.83 128.12.tif

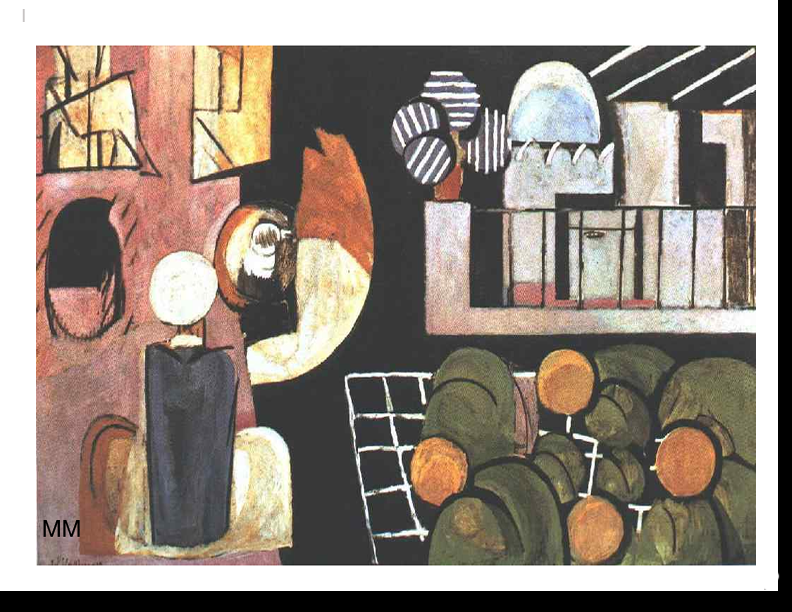

### 08 165.94 162.77 142.83 128.12.tif

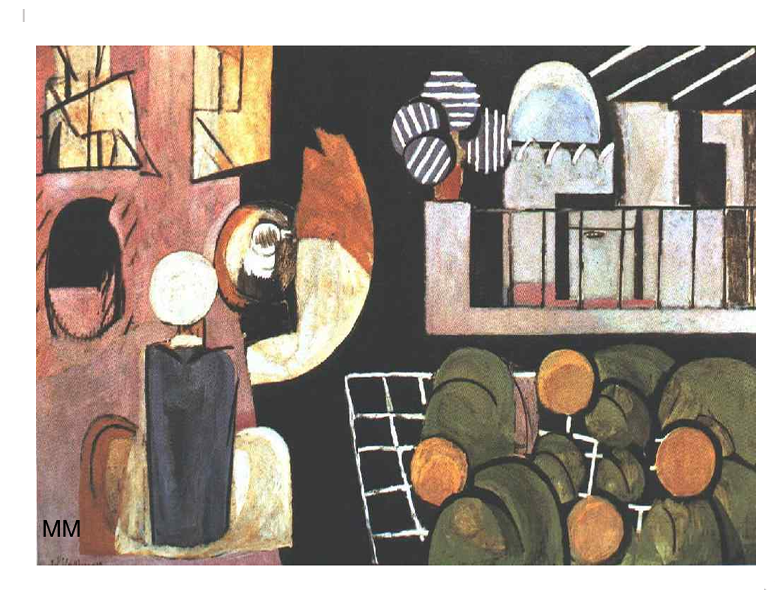

### 08 166.5 164.68 141.13 149.03.tif

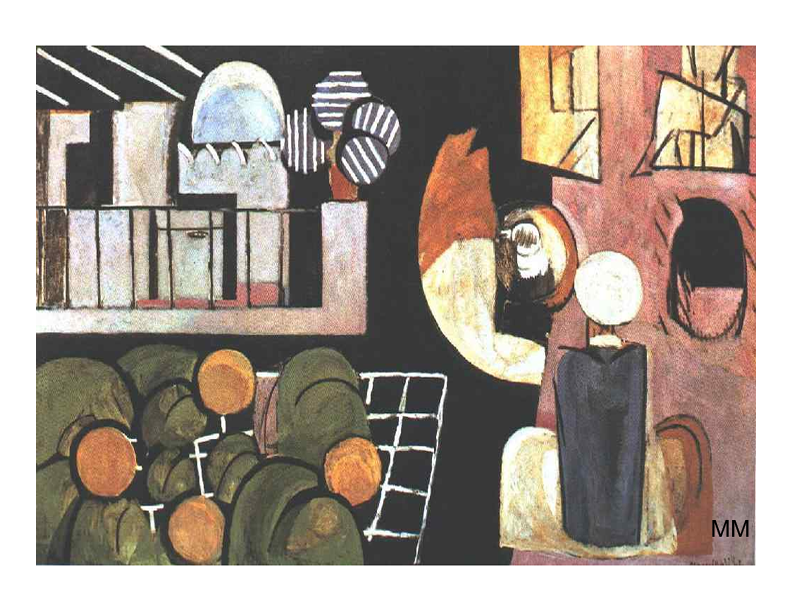

### 09 141.68 144.81 141.68 145.88 589x777 .tif

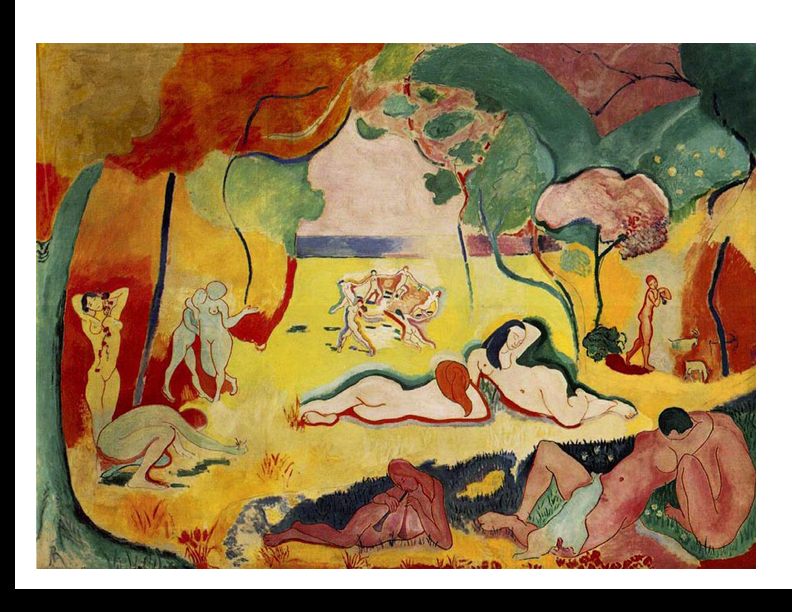

### 09 141.68 144.81 141.68 145.88 589x777 .tif

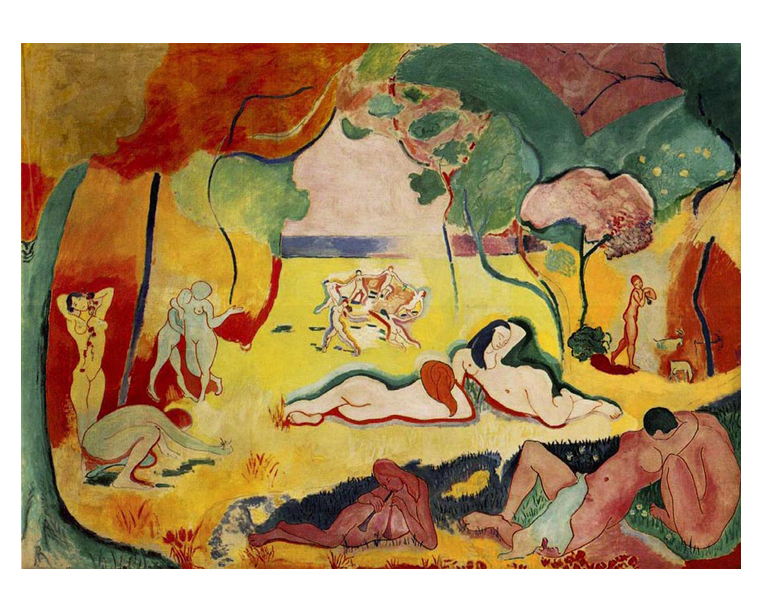

### 09 142.19 149.34 138.51 149.37.tif

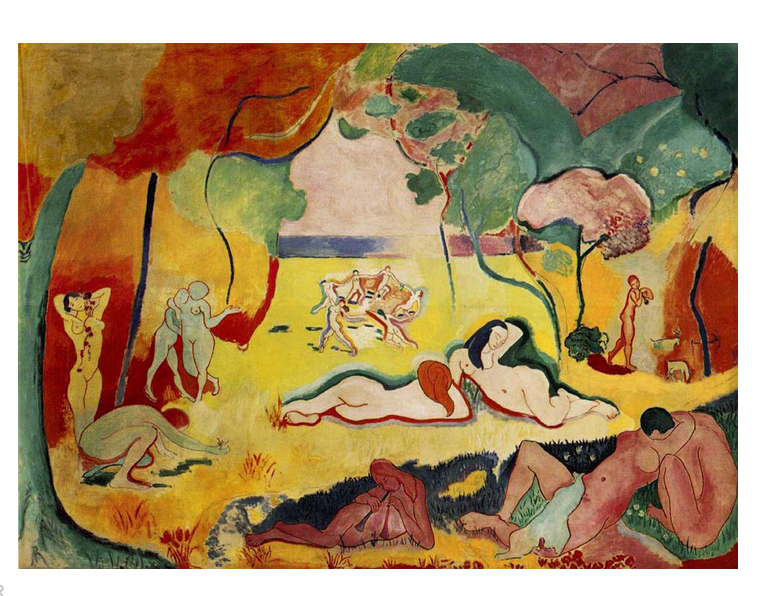

### 09 142.71 149.48 139.13 149.48.tif

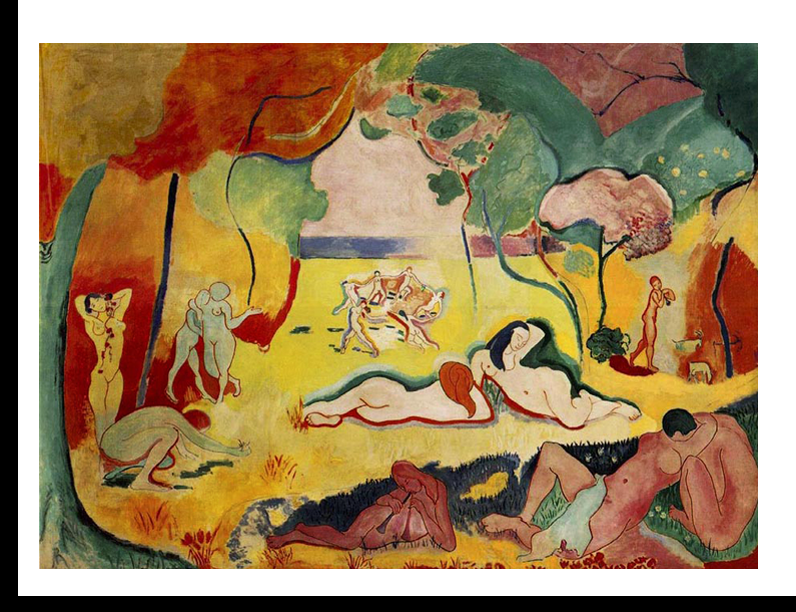

### 10 140.15 140.37 149.36 149.54 1.065.tif

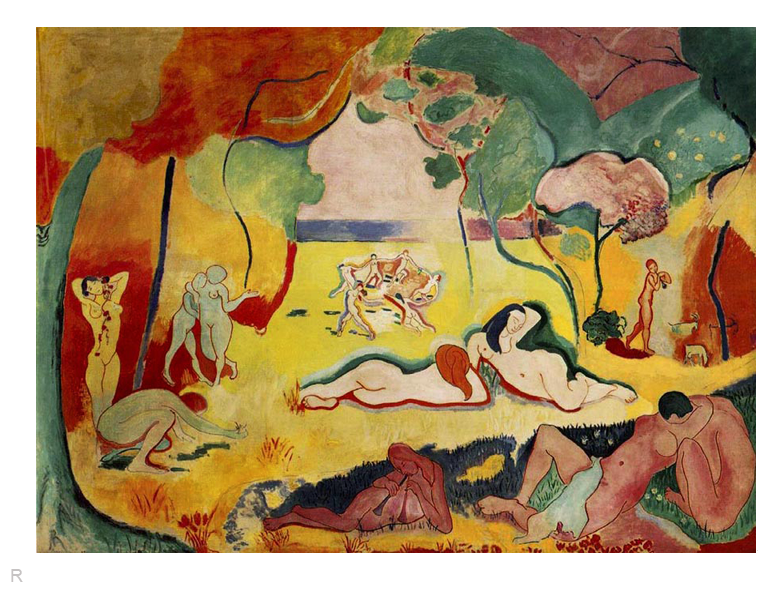

### 10 140.65 140.63 149.79 149.73 1.065.tif

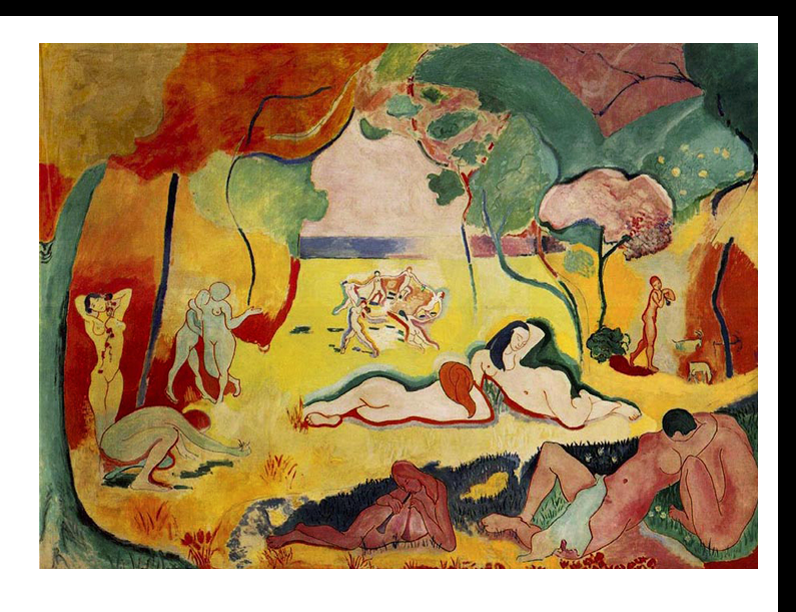

### 10 146.62 146.05 153.89 153.51 1.053.tif

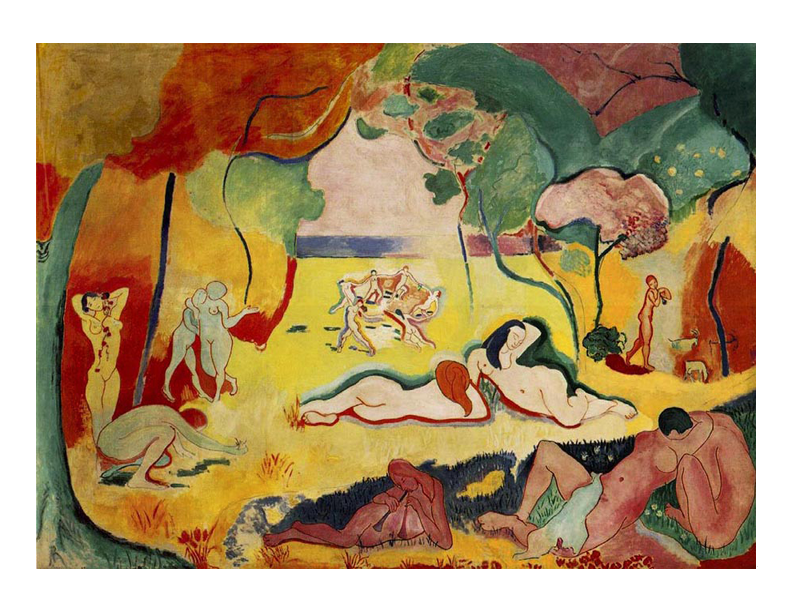

### 11 129.79 126.28 166.32 165.31 601x778 top.tif

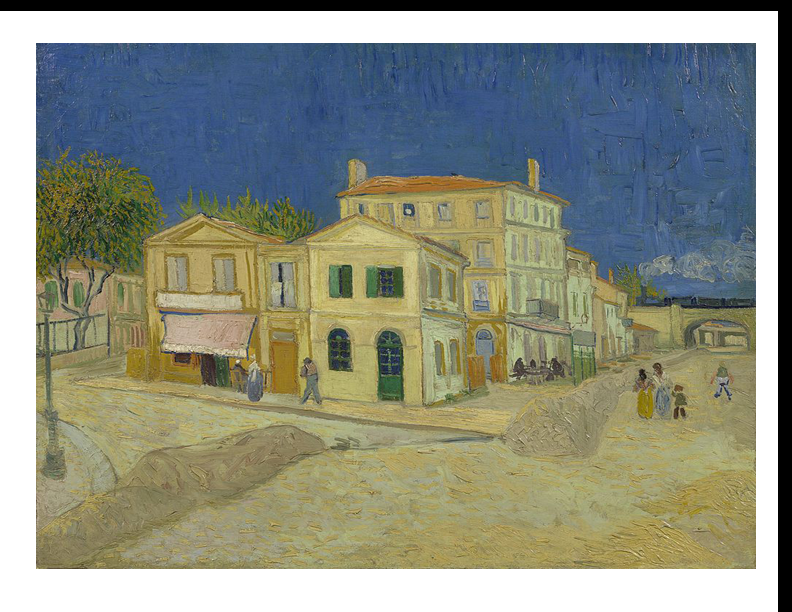

### 11 129.79 126.28 166.32 165.31 601x778 top.tif

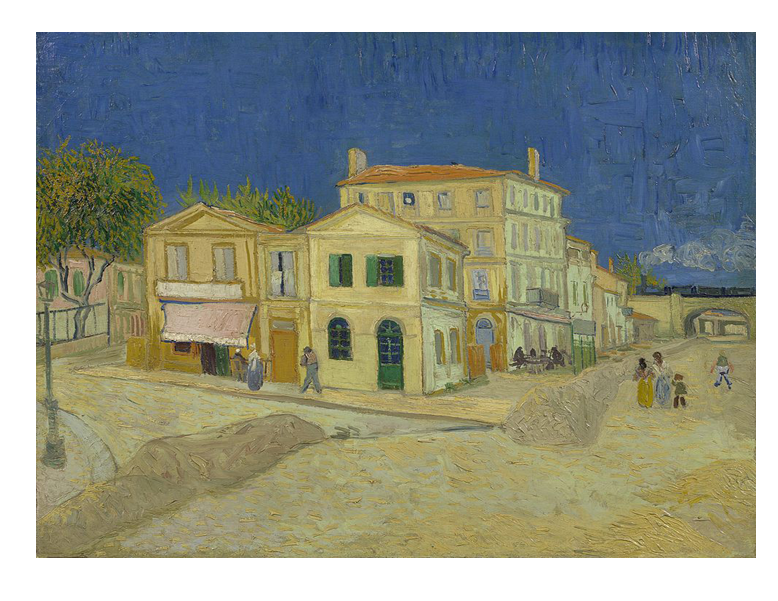

### 11 130.1 124.06 167.33 164.67.tif

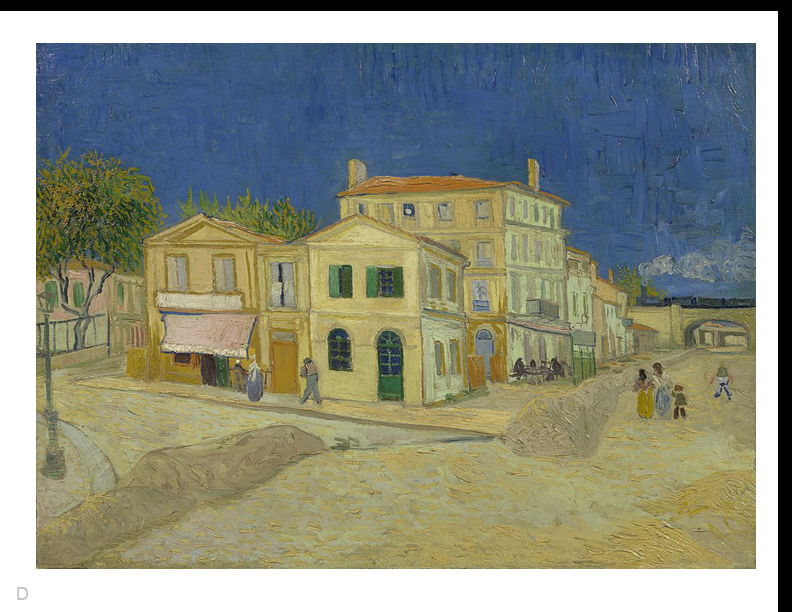

### 11 130.1 124.06 167.33 164.67.tif

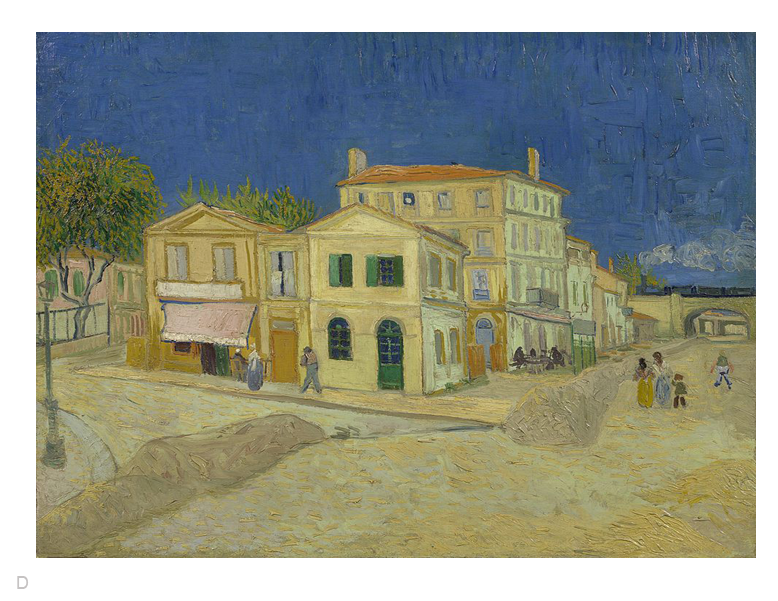

### 12 128.93 133.36 157.5 161.3.tif

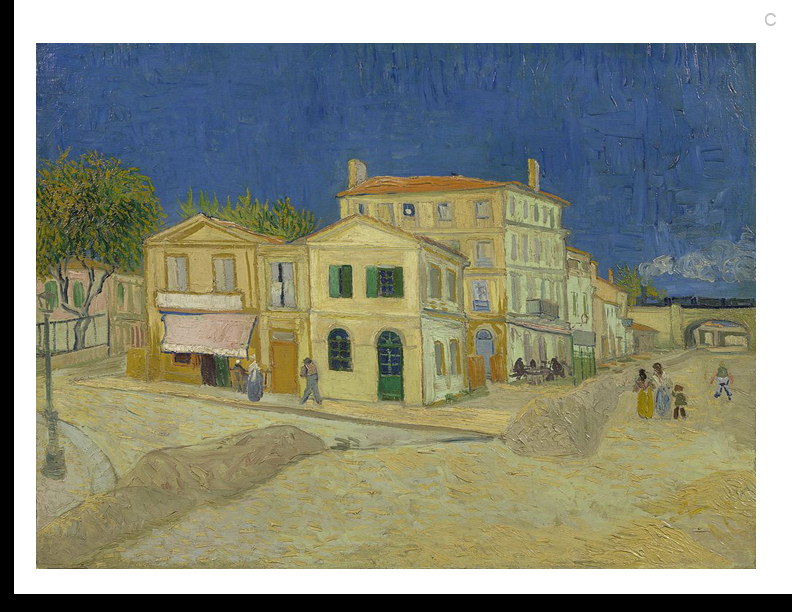

### 12 128.93 133.36 157.5 161.3.tif

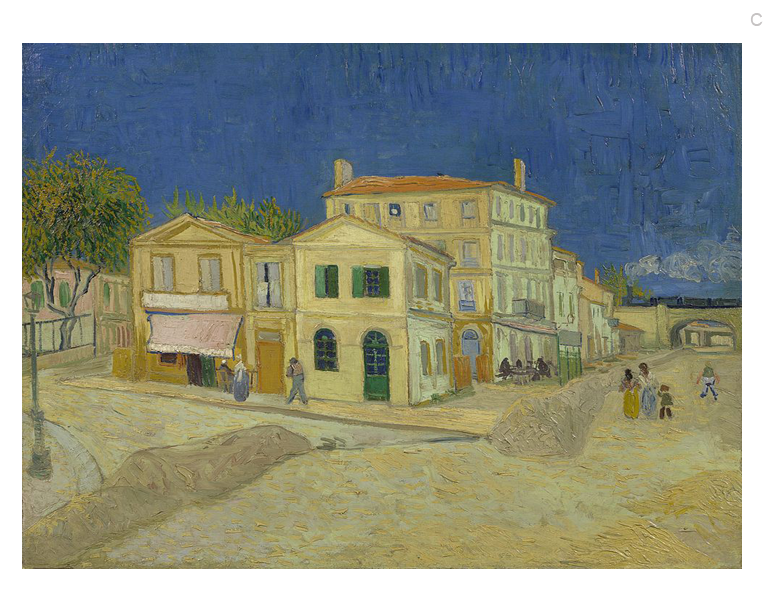

### 12 133.73 135.20 165.82 167.22.tif

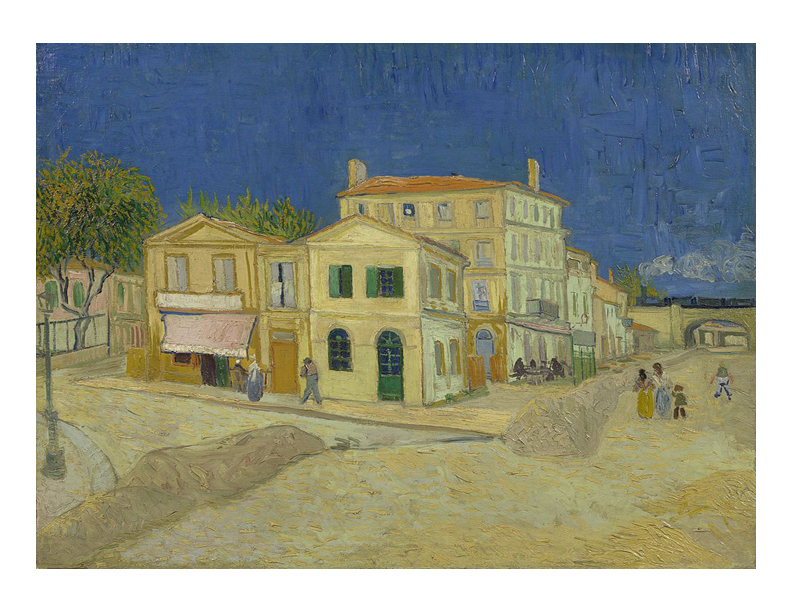

### 13 120.40 120.78 143.56 142.85 584x782 upper.tif

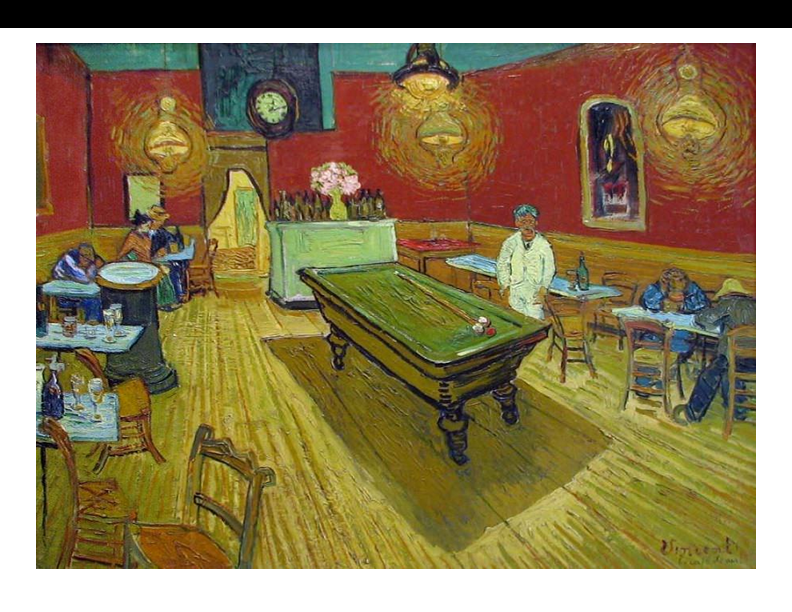

### 13 120.40 120.78 143.56 142.85 584x782 upper.tif

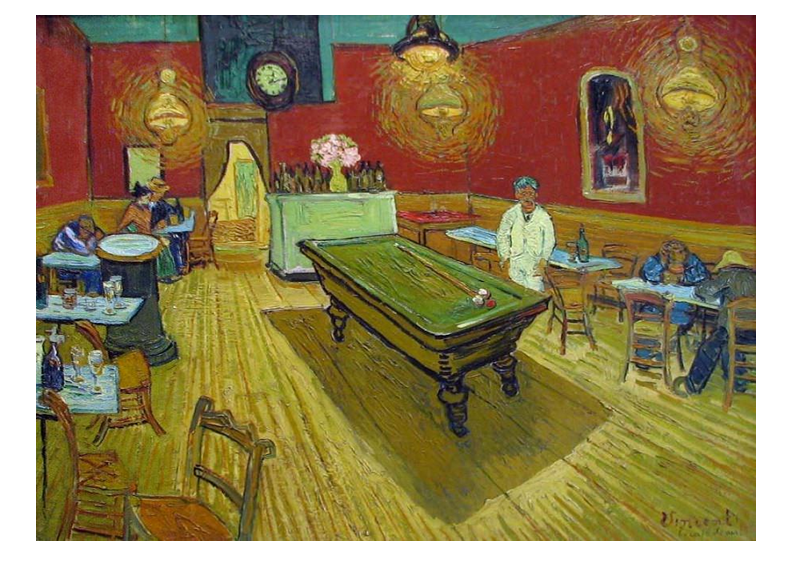

### 13 127.56 138.50 120.19 133.80.tif

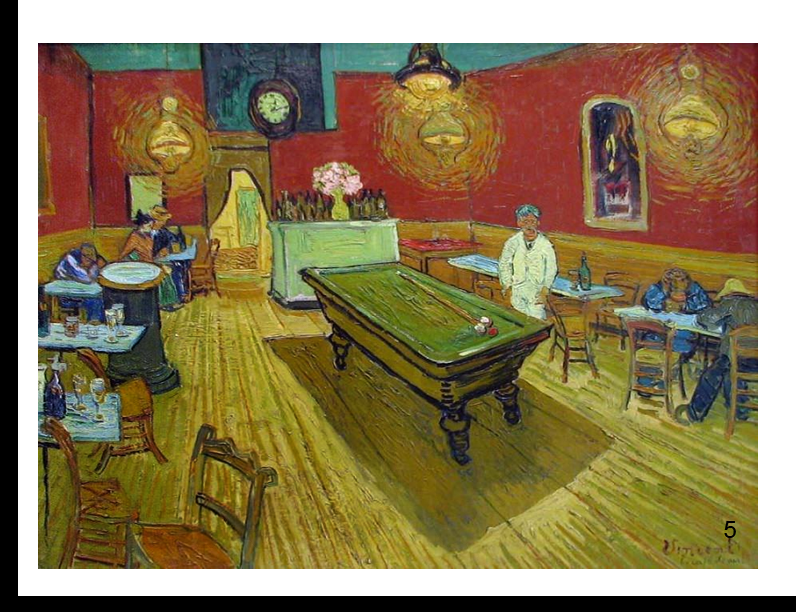

### 13 128.30 137.88 120.63 132.86.tif

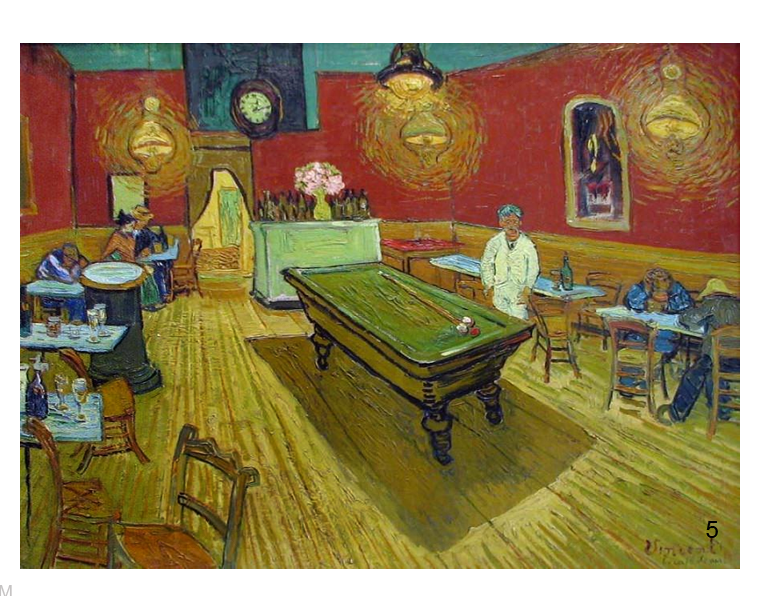

### 14 125.25 125.23 134.76 134.75 1.076..tif

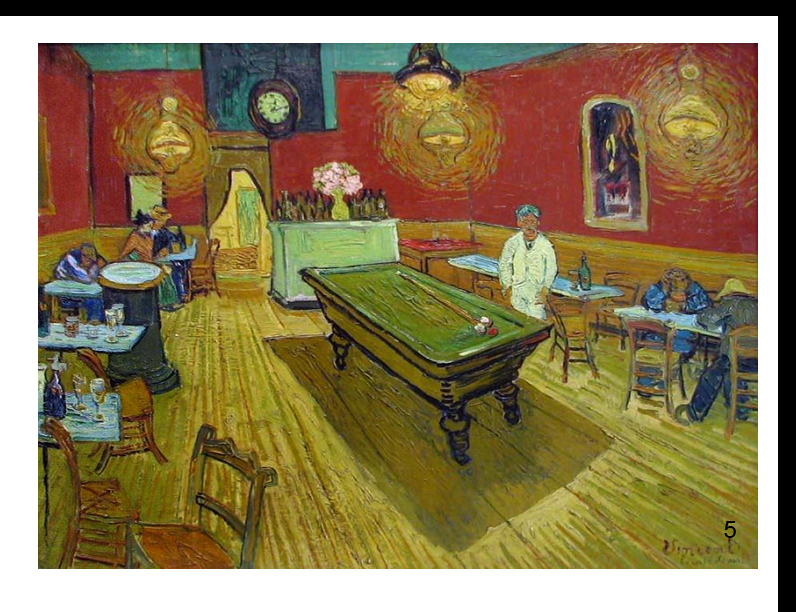

### 14 125.33 125.23 134.52 134.57 1.076.tif

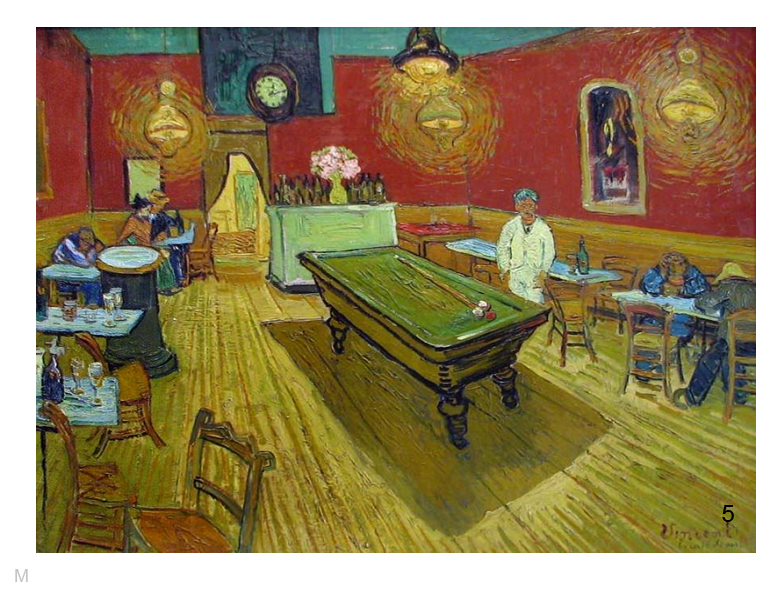

### 14 Cafe 132.99 132.71 142.23 142,19 1.07.tif

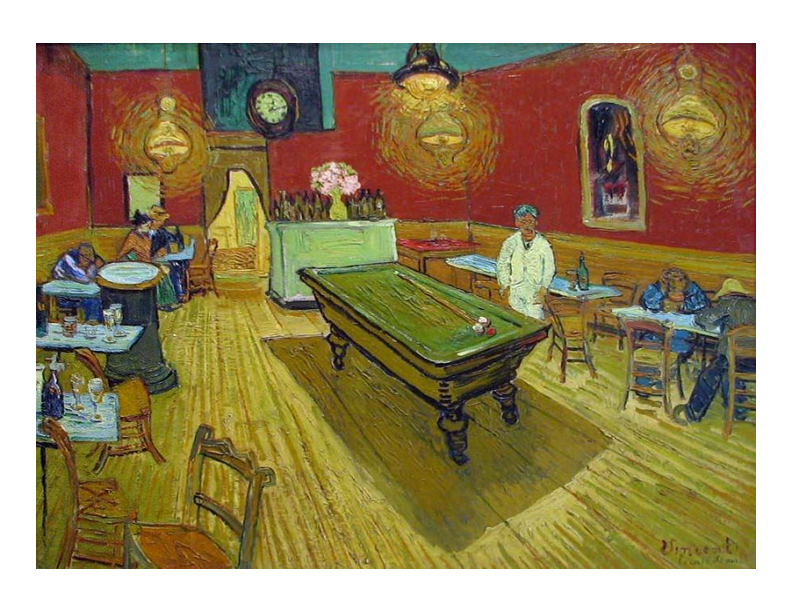

### 15 122.47 122.21 132.27 132.07 1.08.tif

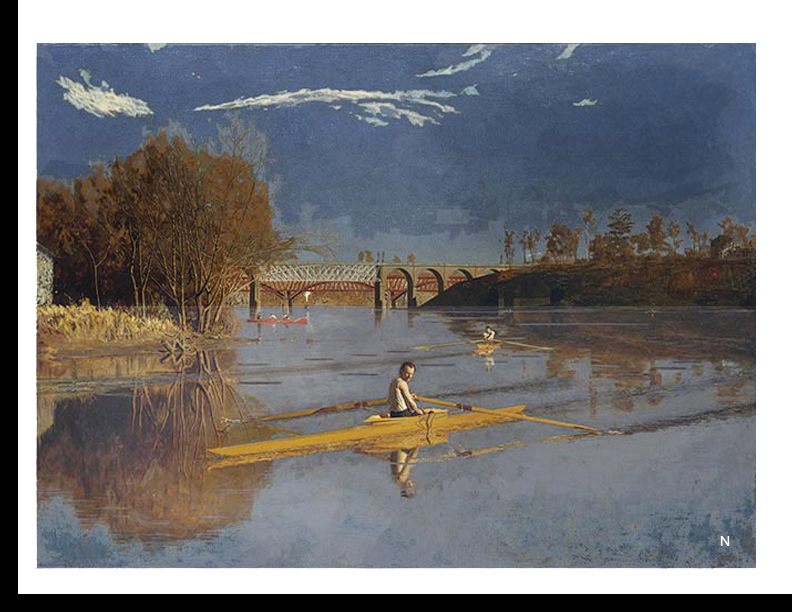

### 15 122.47 122.21 132.27 132.07 1.08.tif

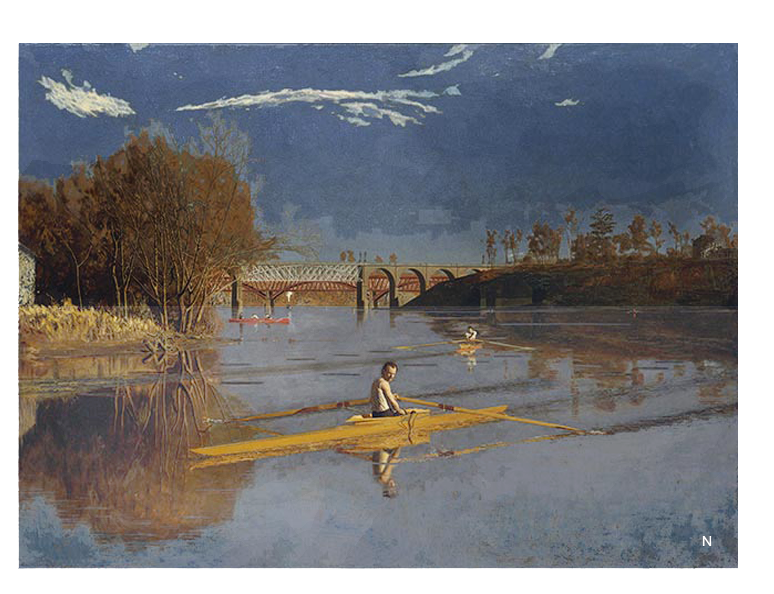

### 15 129.18 129.13 138.57 138.58 1.073.tif

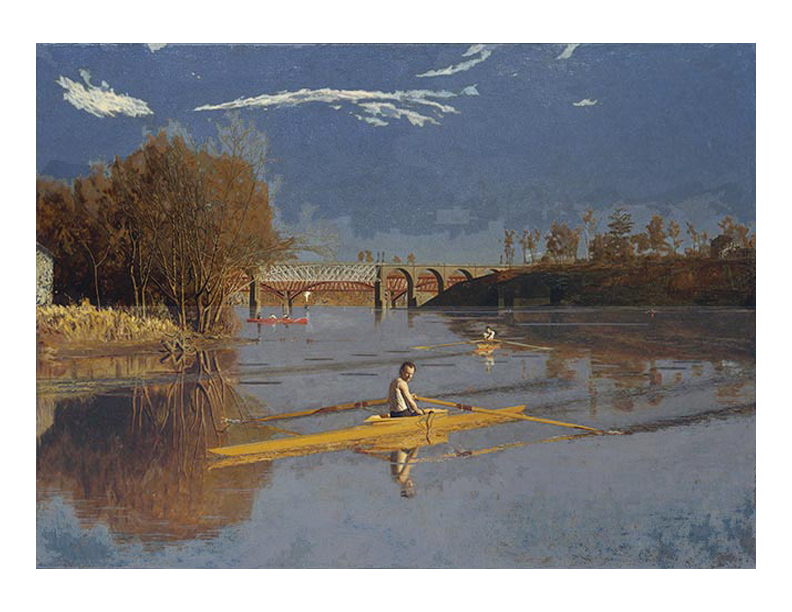

### 16 114.89 120.78 133.15 140..11 595 x 772 upper.tif

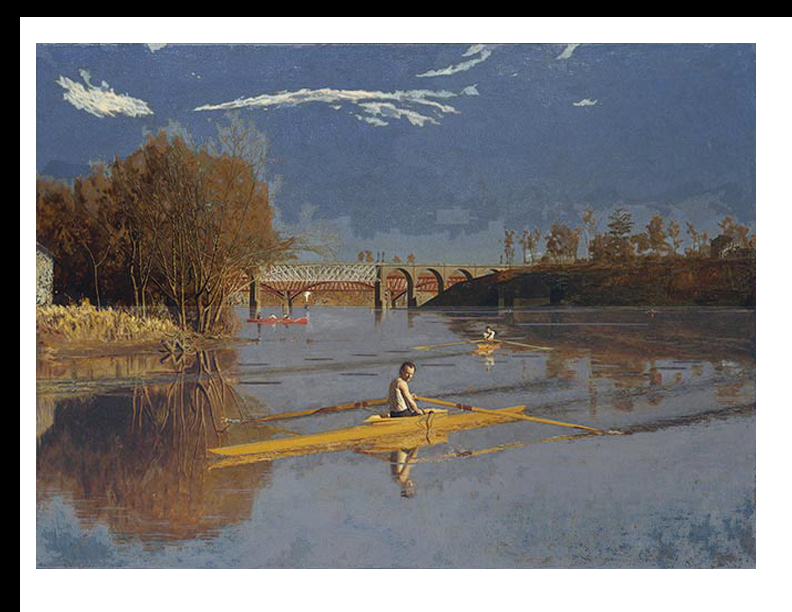

### 16 114.89 120.78 133.15 140..11 595 x 772 upper.tif

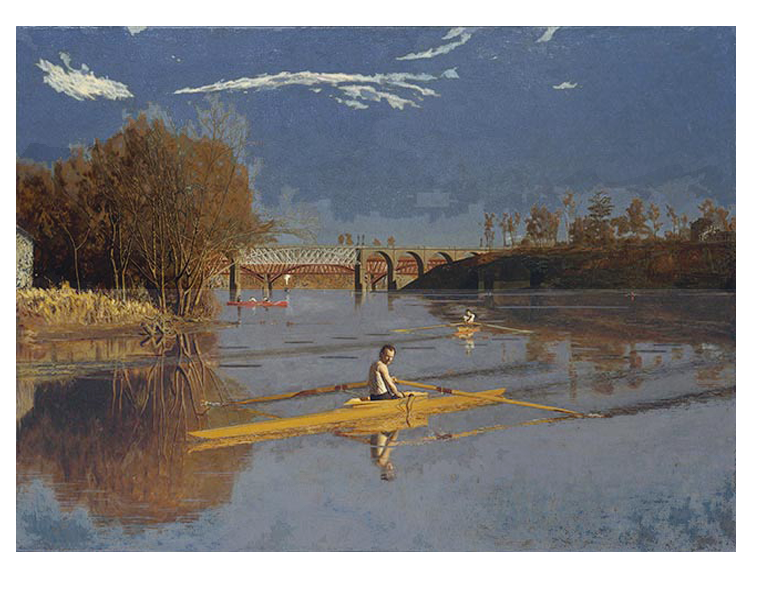

### 16 119.77 105.43 146.55 137.69.tif

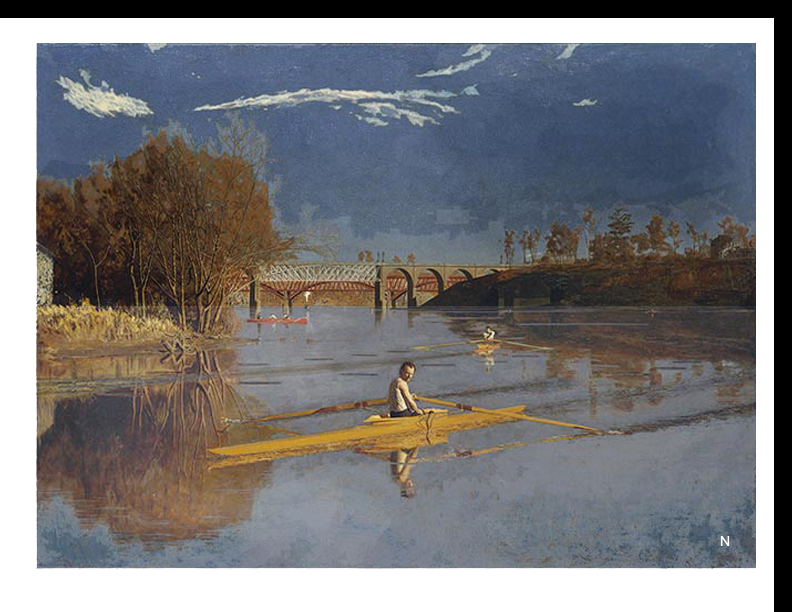

### 16 119.77 105.43 146.55 137.69.tif

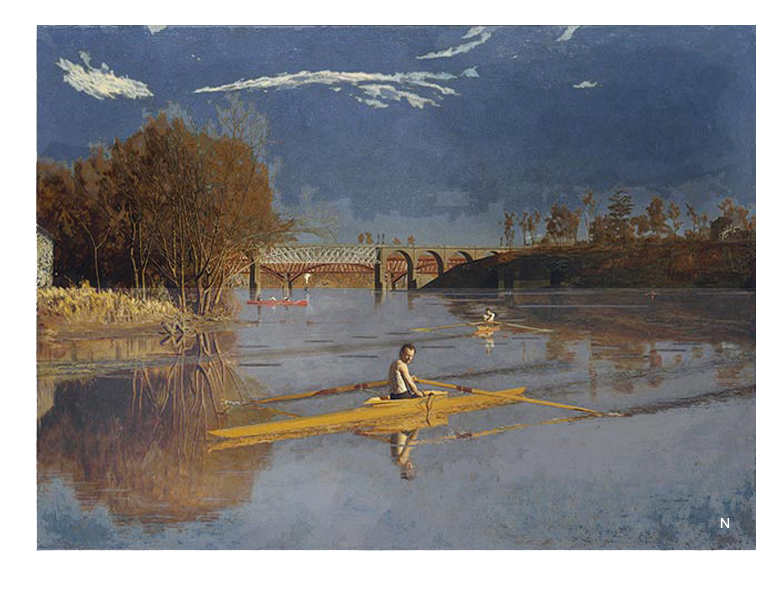

### 17 147.03 133.18 121.83 121.77.tif

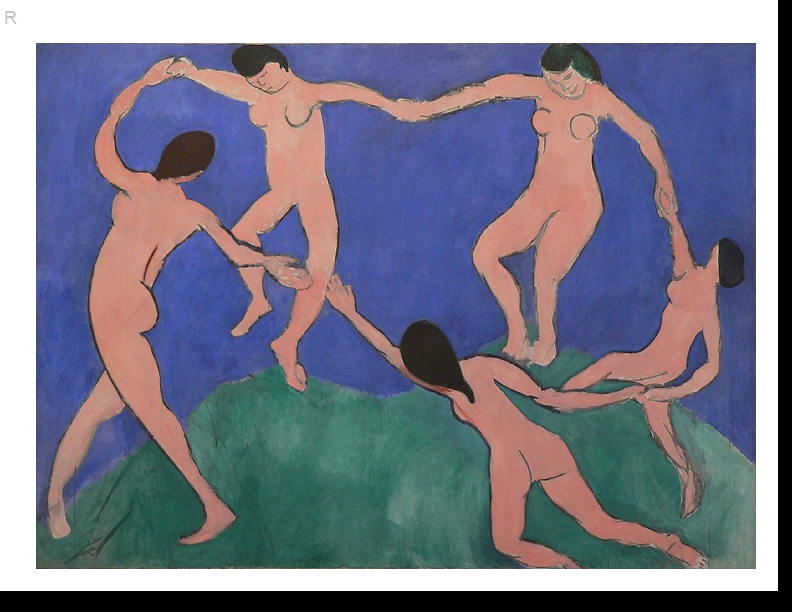

### 17 147.03 133.18 121.83 121.77.tif

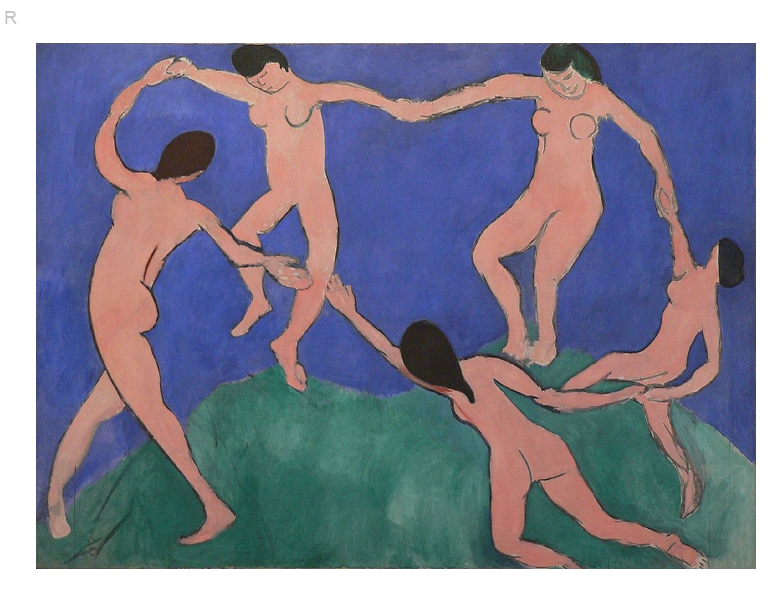

### 17 Dance 145.75 136.82 130.68 135.96.tif

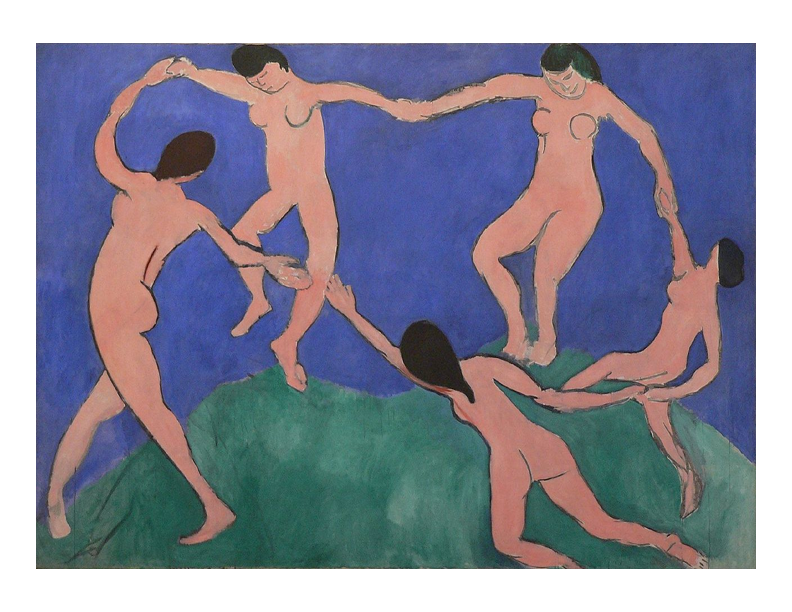

### 18 Dance 132.49 127.88 126.35 137.12 591x778 upper.tif

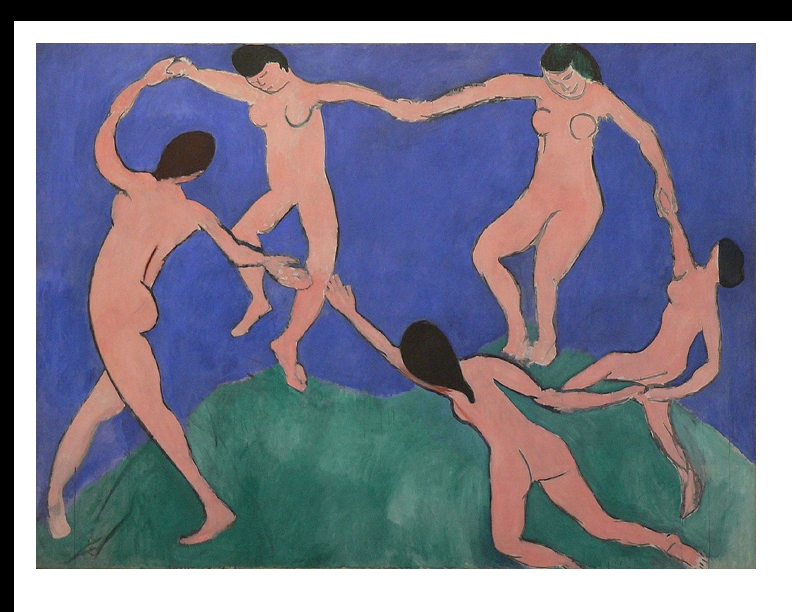

### 18 Dance 132.49 127.88 126.35 137.12 591x778 upper.tif

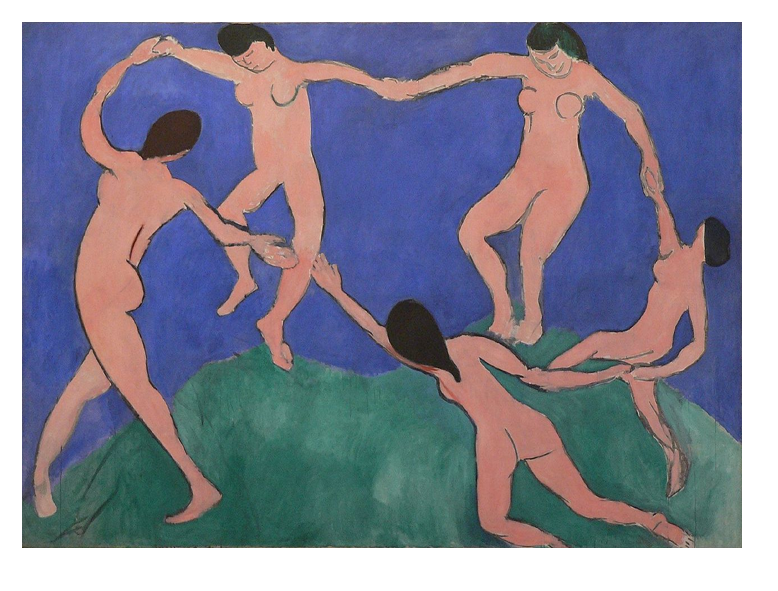

### 18 Dance 132.49 127.88 126.35 137.12.tif

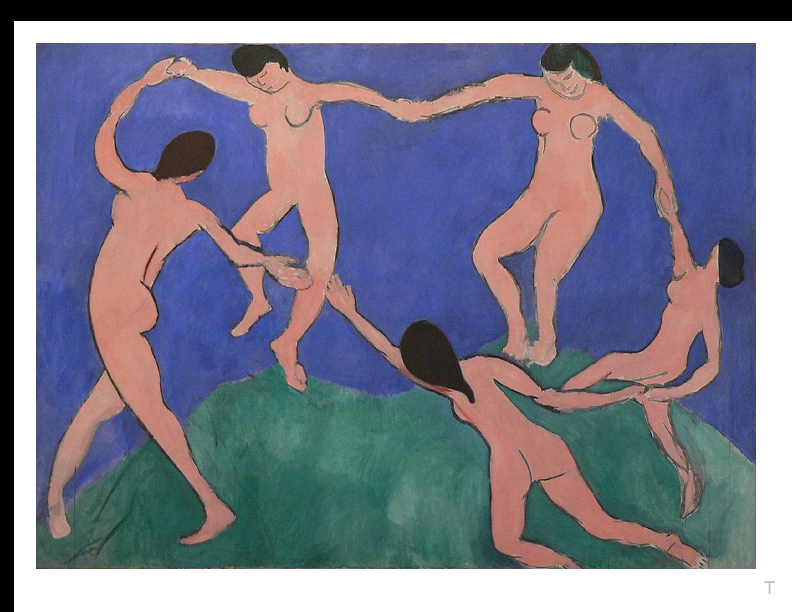

### 18 Dance 132.49 127.88 126.35 137.12.tif

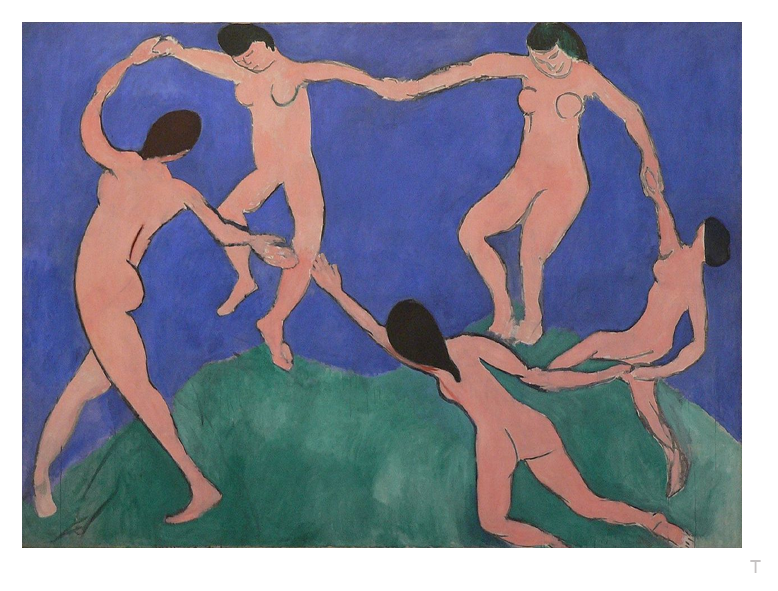

### 19 119.02 138.55 104.37 141.96.tif

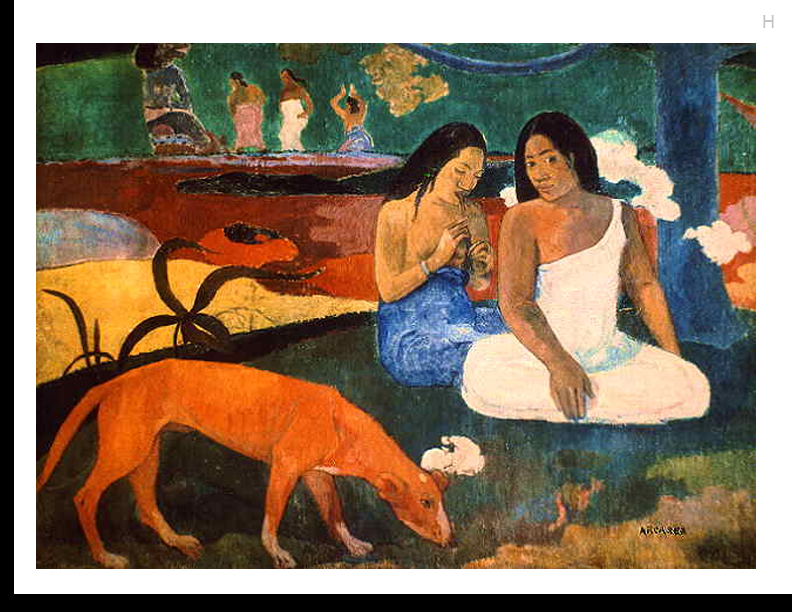

### 19 119.02 138.55 104.37 141.96.tif

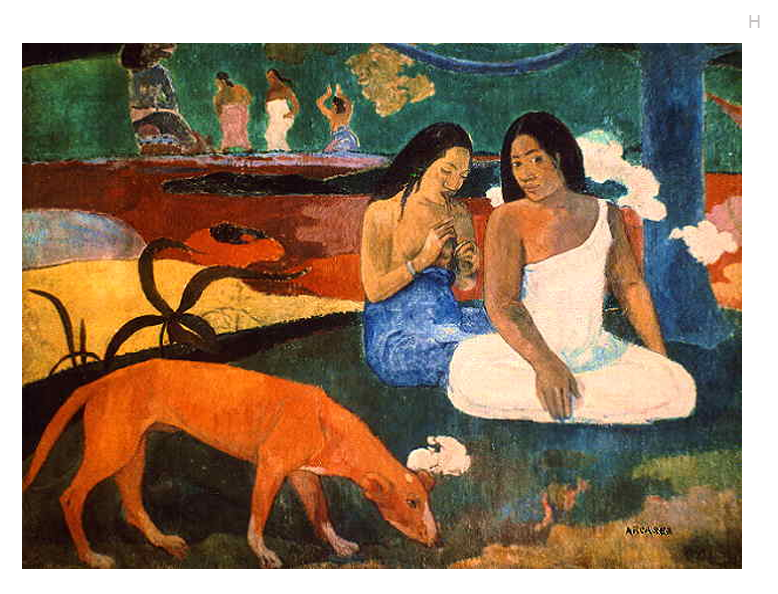

### 19 123.12 134.10 109.01 134.89 591x777 lower.tif

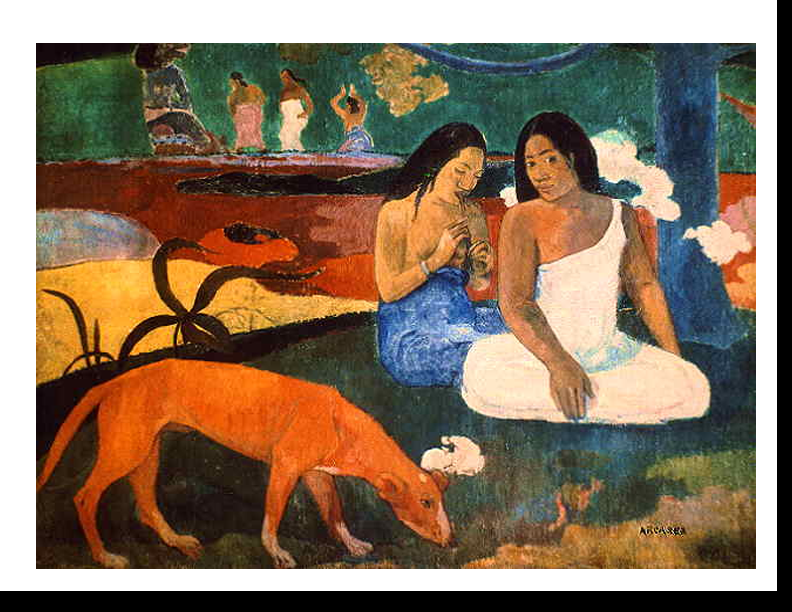

### 19 123.12 134.10 109.01 134.89 591x777 lower.tif

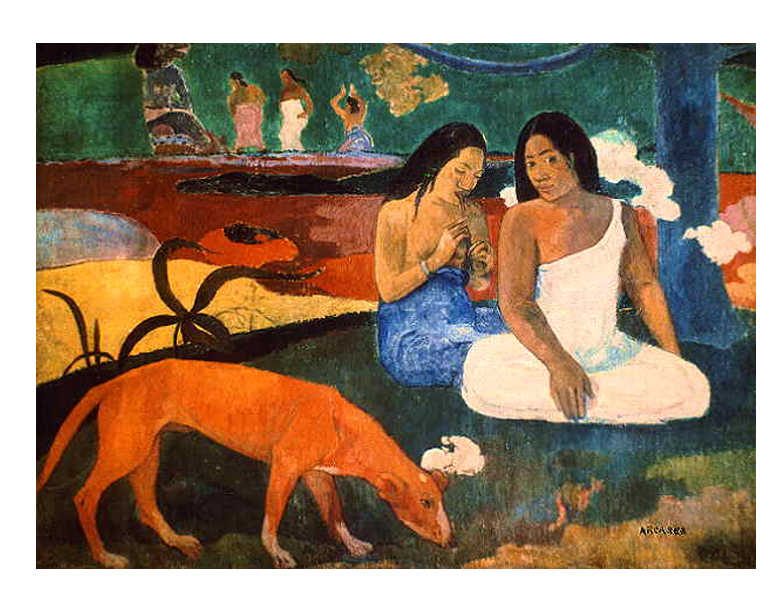

### 20 117.76 127.27 115.96 143.58.tif

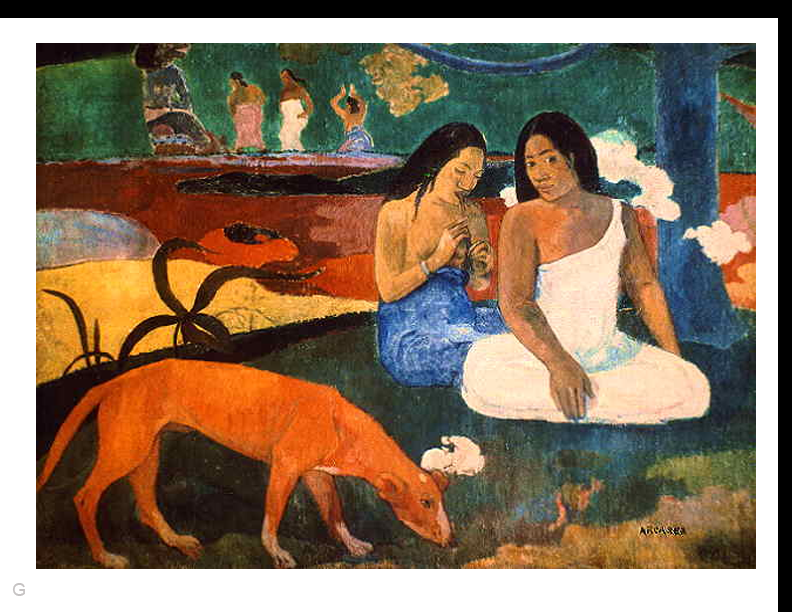

### 20 117.76 127.27 115.96 143.58.tif

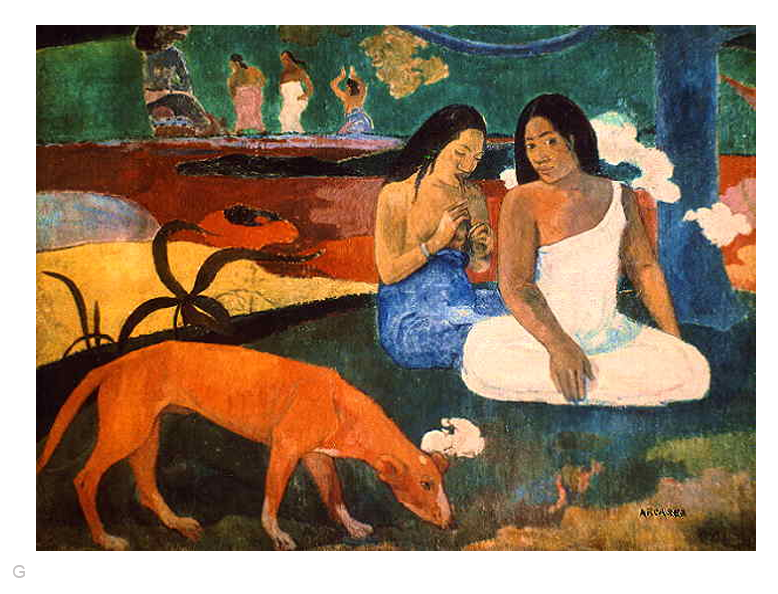

### 20 124.87 138.70 117.01 147.92.tif

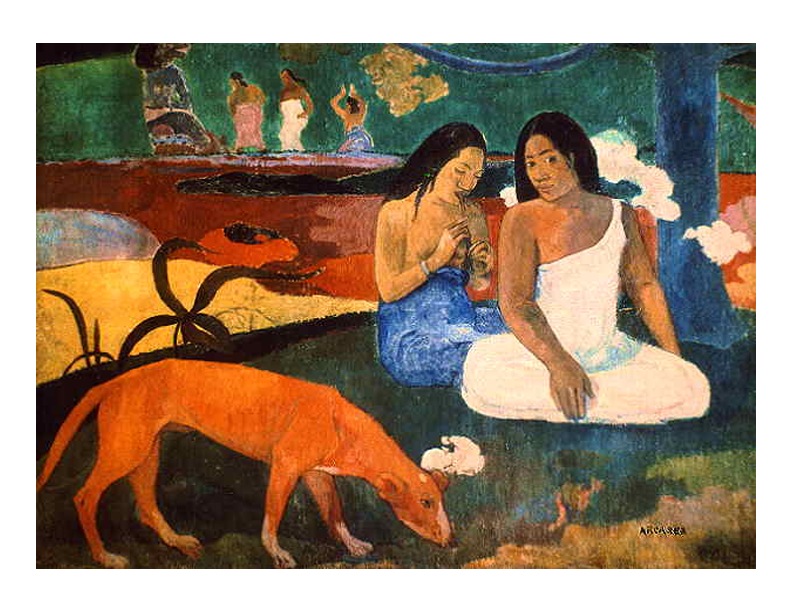

### Figure 3 82.63 82.56 124.73 124.77.tif

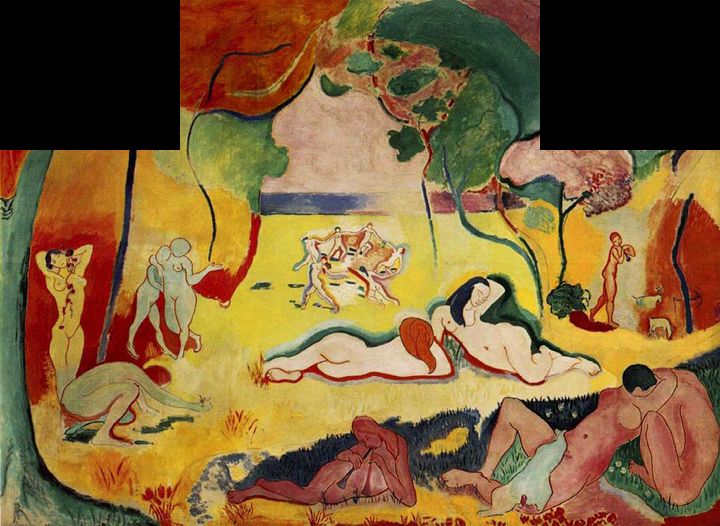

### Figure 4 115.52 115.56 124.73 124.80 1.08.tif

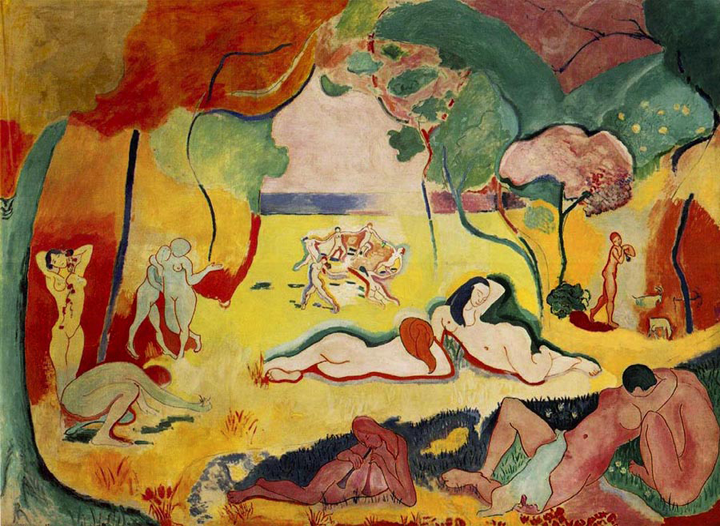
